# Living by the sea: chromosome-scale genome assembly and salt gland transcriptomes provide insights into ion regulatory mechanisms in the saline-tolerant mosquito *Aedes togoi*

**DOI:** 10.64898/2026.04.09.717544

**Authors:** Jonathan Chiang, Elissa Khodikian, Orna Phelan, Ana K. Parra, Daniel A. H. Peach, Andrea C. Durant, Benjamin J. Matthews

## Abstract

The coastal rock pool mosquito, *Aedes togoi*, is among the few saline-tolerant mosquito species who lay their eggs in seawater pools where their larvae develop in water that spans dilute freshwater to hypersaline conditions. *Ae. togoi* is found in a relatively restricted range spanning the North Pacific coast of North America and coastal regions of Asia from subtropical to subarctic latitudes. Here, we present a *de-novo* chromosome-scale genome assembly and gene annotation for *Ae. togoi*, highlighting its relatively small genome size and novel chromosomal arrangements compared to other available genomes of *Aedine* mosquitoes. As part of the annotation process, we detail repeat content and distribution and curate several key multi-gene families, focusing on ion-transport proteins enriched in the larval salt-secreting gland that are candidates for facilitating hyperosmotic urine formation during development in saline water. Using these new resources, we gain mechanistic insight into the ion regulatory capabilities that power the remarkable saline tolerance of the larvae of *Ae. togoi*. Altogether, we have contributed to the growing body of genomic and transcriptomic resources for diverse mosquito species and provided mechanistic insights into the molecular adaptations required for an insect to thrive in highly dynamic environments such as coastal rock pools.

## Introduction

Recent advances in long-read sequencing and chromatin conformation capture technologies have revolutionized our ability to study mosquito genomes in their full complexity. Previous high-quality, chromosome-scale assemblies have revealed striking differences in repetitive element content, genome organization, and chromosome structure, together providing a greater understanding of mosquito evolution (Derilus et al., 2025; Habtewold et al., 2024; Hesson et al., 2025; Liu et al., 2023; Matthews et al., 2018; Morinaga et al., 2025; Ryazansky et al., 2024a; Zamyatin et al., 2021). These resources have paved the way for fundamental discoveries into many processes of mosquito behaviour and physiology. With direct implications on mosquito-vectored disease control strategies, these resources have allowed for targeted characterization of the genes and molecular mechanisms underpinning vector competence, vector transmission, and environmental adaptation.

Mosquitoes are known to be highly adaptable to environmental conditions and stressors, leading to their global distribution and establishment in almost every habitat with standing water available, notably in both rural and urban environments. Adapting to such a wide range of environments requires the adoption of suitable anatomical, physiological, and behavioural strategies. The mechanisms by which mosquitoes adapt to new or changing environments has implications for the habitats and geographical regions they are able to access (and potentially serve as vectors for disease), both currently and in the future as the environmental characteristics of a region shifts. Further research into the genetic underpinnings of climate tolerance and water composition tolerance, two major factors deciding habitat suitability, will provide a greater understanding and predictive ability of the geographic range of a mosquito species (Ghosh et al., 2020). We propose the coastal rockpool mosquito, *Aedes togoi*, as a study species well-suited for the characterization of cold-weather and water-salinity tolerances of mosquitoes across the life cycle.

*Ae. togoi* is found along the Northern Pacific coastline of North America and in coastal regions of East Asia, spanning subtropical to subarctic latitudes, where it is a potential vector for flaviviruses and filarial parasites (Peach and Matthews, 2020; Rosen et al., 1978; Tanaka et al., 1979). Like other mosquito species, *Ae. togoi* require a suitable water source to continue their life cycle: a place to lay their eggs and for their offspring to develop in. Larvae of most mosquito species are restricted to development in clean freshwater; however, *Ae. togoi* are among the 5% of euryhaline species that can survive and develop in brackish and saline waters (Asakura, 1980; Bradley, 1987; Choi and Choi, 2024; McGinnis and Brust, 1983; Trimble and Wellington, 1979, Patrick and Bradley, 2000). Inhabiting coastline regions may have been an adaptation to occupy an ecological niche with less competition with freshwater-obligate species (Lushasi et al., 2024; Noden et al., 2015). The coastal rock pools that *Ae. togoi* inhabit experience wide ranging fluctuations in environmental conditions, most notably temperature and salinity, as they are generally unshaded and filled by both salt spray and rainfall.

In freshwater, hypo-osmotic conditions pose an osmo-regulatory need for aquatic animals to counteract passive ion loss and water gain. On the other hand, in hyper-osmotic conditions, such as in seawater, there is an increased risk of desiccation to the ion-concentrated surrounding environment. Euryhaline mosquito larvae that develop in coastal habitats regularly experience rapid changes in water salinities ranging from dilute to hypersaline several times more concentrated than seawater, and *Ae. togoi* are no exception (Asakura, 1980; Phelan, 2023). The physiological feats of euryhaline mosquito larvae during their development in these habitats of salinity extremes are indeed impressive; a larva maintains a near-constant hemolymph osmotic pressure by hyper-regulating in dilute freshwater by actively absorbing ions and hypo-regulating in saline water by actively excretingions (Bradley and Phillips, 1977; Asakura, 1980; Bradley, 1987; Donini et al., 2005). The physiological mechanisms to accomplish this are partially resolved for some species (Patrick et al., 2024, Smith et al., 2008) but are almost entirely unknown for *Ae. togoi*.

In other euryhaline mosquitoes, saline conditions have selected for anatomical modifications of the larval hindgut that involves specialized segments that actively secrete salts and concentrate the urine. Two examples that span mosquito genera include the posterior rectal segment of saline-tolerant larvae of *Aedes* species (Patrick et al., 2024; Albers and Bradley, 2011), and the specialized salt-secreting rectal cells in *Anopheles* species (Smith et al., 2008). Additionally, previous characterization of proteins involved in ion regulation in euryhaline species has documented how expression of ion-pump proteins can adapt to changes in osmotic pressure and osmoregulatory needs (Patrick et al., 2024; Smith et al., 2008). Instead of being situated near or within the rectum, the specialized salt-secreting organ of *Ae. togoi larvae* has emerged further posteriorly along the hindgut, as an elaboration of part of the anal canal (Asakura, 1980; Asakura, 1982).

Knowledge of the ion-transport machinery that drives the production of a hyperosmotic urine via the salt-secreting gland of euryhaline larvae is very much in its infancy. However, recent work on *Aedes taeniorhynchus* larvae demonstrates that active sodium secretion is powered by the V-type H^+^-ATPase. Does the salt-secreting gland of *Ae. togoi* larvae employ a similar strategy to achieve ion balance? To address this question and other aspects of *Ae. togoi’s* unique biology, we present a *de-novo* chromosome-scale genome assembly and accompanying protein-coding gene-set annotations. Furthermore, we use a bulk RNA-sequencing approach from both whole larval tissues and salt-secreting gland tissues from animals reared in freshwater (FW) or seawater (SW) to begin to resolve ion regulatory pathways and highlight candidate transcripts that are potentially important to these processes. Together, these data and resources contribute useful reference points to support future comparative physiology and evolutionary genomics studies, provide key insight into osmoregulation in euryhaline mosquitoes, and lay a solid foundation for further physiological studies in *Aedes togoi*.

## Results

### Generation of an *Ae. togoi* colony from British Columbia, Canada

To facilitate the generation of a reference genome for *Aedes togoi*, we first established a laboratory colony from field collections in Lighthouse Park, West Vancouver, Canada (Figure 1A). In addition to collecting larvae and pupae, we recorded water temperature, salinity, and larval and pupal abundance data from coastal rock pools over the course of a year to define the field conditions that this population of *Ae. togoi* might routinely encounter and tolerate (Supplementary Table 3). The large majority of *Ae. togoi* larvae and pupae were found during the summer months (May-September), when water temperature and salinity were higher (Supplementary Table 3). These weekly data showcase how this population of *Ae. togoi* larvae and pupae can tolerate water temperature from ∼1.5-30°C and salinity from 0-84.8 ppt (Figure 1B).

**Figure 1.**
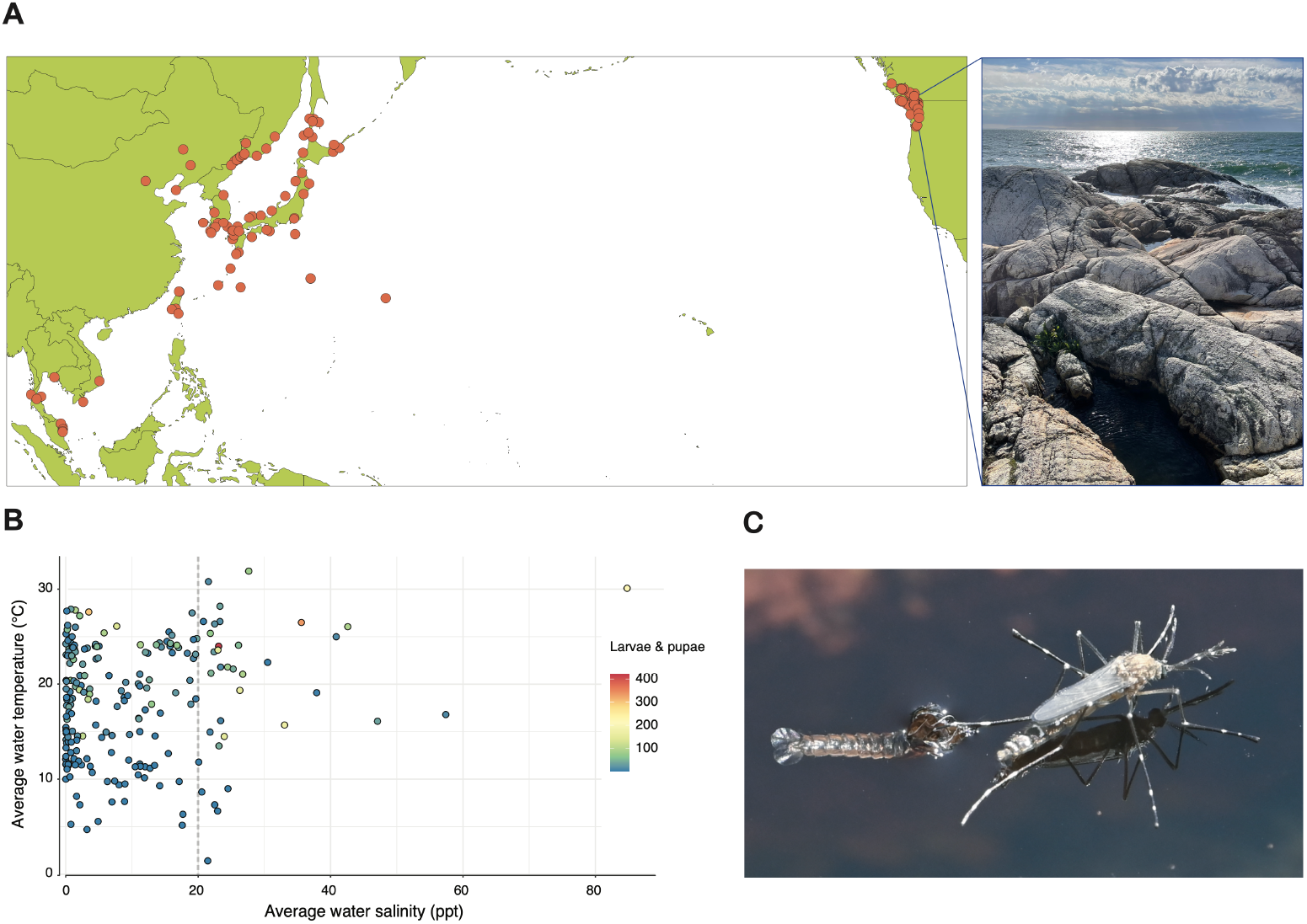
Salinity tolerance and global distribution of *Aedes togoi*. **(A)** Global distribution of *Ae. togoi*, with locations where *Ae. togoi* are present indicated in orange. Observation data re-plotted from Peach and Matthews, 2020. Right side-panel shows *Ae. togoi* coastal rockpool habitat in Lighthouse Park in West Vancouver, Canada. **(B)** Water temperature (°C) and salinity (ppt) of coastal rockpools with *Ae. togoi* larvae or pupae present were measured over the course of a year to capture the range of conditions that *Ae. togoi* are exposed to. The dotted-line at 20 ppt represents the average surface water salinity of seawater in the area surrounding the sampling site. Colour represents the number of *Ae. togoi* larvae and pupae sampled in a rockpool on a single day. **(C)** Adult *Ae. togoi* newly emerged from coastal rockpool.

### DNA and RNA sequencing

As the basis for *de novo* reference genome assembly, we extracted high molecular weight (HMW) genomic DNA from *Ae. togoi* pupae taken from our laboratory colony. HMW DNA collected from a single male pupa was used to generate 4.25 million Q20 HiFi reads across two flow-cells, with a combined mean read length of 13.4kb, representing 56.8 Gbp sequenced. From another single male pupa, 68.3 Gbp of Hi-C data were collected yielding 424.6 million paired-end Illumina reads. Additionally, 17.5Gbp across 2.5 million Nanopore (ONT) reads with were generated from a single male pupa, 12.2 Gbp across 2.1 million ONT reads were generated from a single female pupa, 23.1 Gbp across 2.1 million ONT reads were generated from a pool of eight male pupae, and 20.4 Gbp across 1.6 million ONT reads were collected from a pool of eight female pupae (Supplementary Table 1).

*Ae. togoi* mRNA datasets were collected from various tissue and life-stages. mRNA collected from *Ae. togoi* adults (Figure 2A) generated 1,680,724,872 raw reads from pooled head libraries and 1,602,957,326 raw reads from pooled body libraries (Supplementary Table 1). Additionally, mRNA extracted from pooled whole larvae generated 182,309,635 raw reads and mRNA from pooled larval salt glands generated 694,868,370 raw reads (Supplementary Table 1).

**Figure 2.**
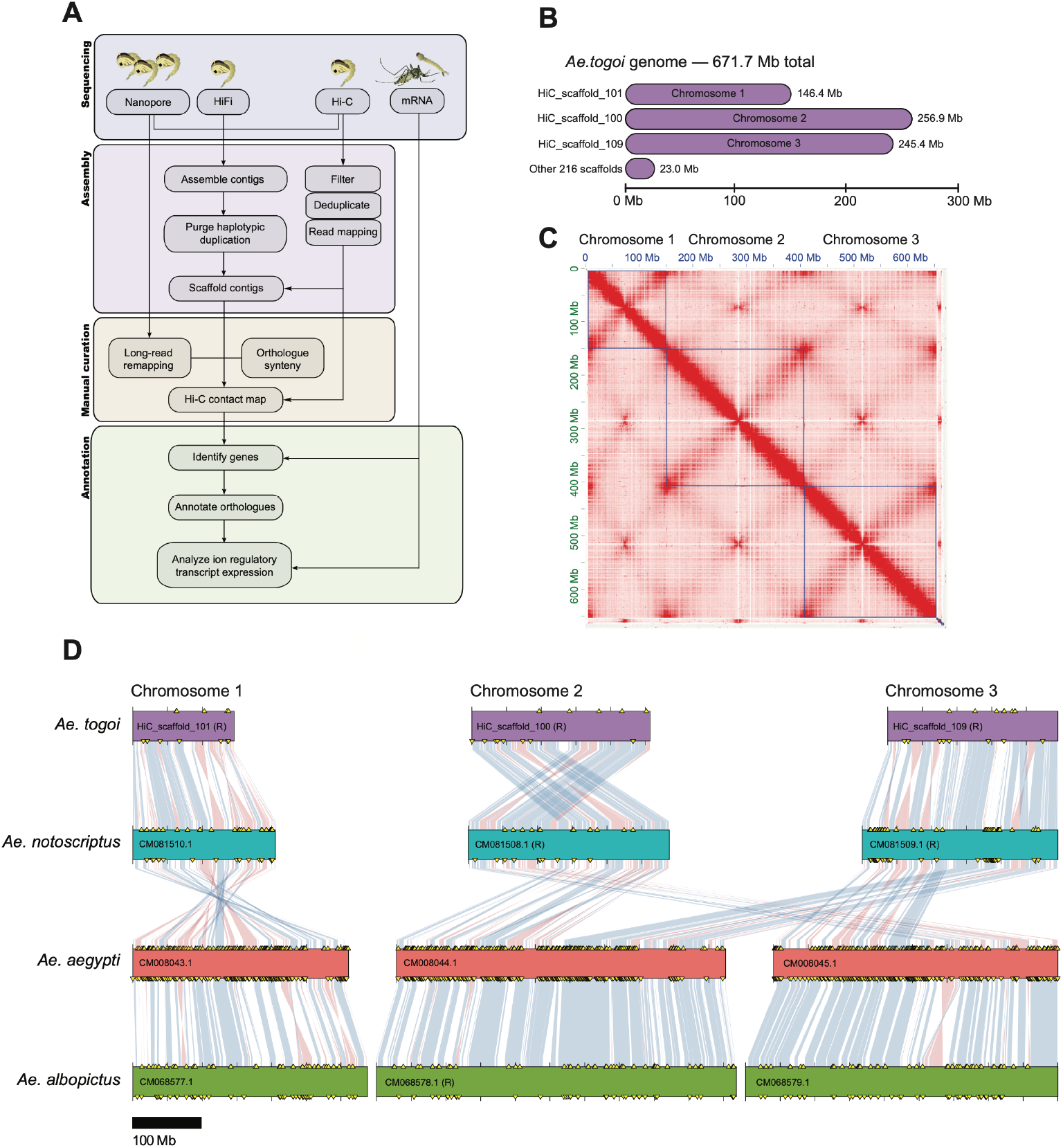
Genome assembly and chromosome structure of the saline-tolerant *Aedes togoi*. **(A)** Overview of genomic and transcriptomic sequencing data collection, genome assembly steps, and gene annotation strategy. **(B)** Scaffold and full genome lengths for the chromosome-scale *Ae. togoi* assembly. **(C)** Hi-C contact map highlighting chromosome structure of the curated *Ae. togoi* genome assembly. **(D)** Chromosomal synteny between *Ae. togoi, Ae. notoscriptus, Ae. aegypti*, and *Ae. albopictus* genomes based on the location of orthologous BUSCO genes. Chromosomal inversions are highlighted in red while syntenic blocks with matching orientation are denoted in blue. Chromosomes with the majority of synteny on the opposite strand are labelled with (R).

### Genome assembly

The initial assembly of HiFi reads, along with Hi-C and ultra-long ONT read integration, resulted in a primary contig assembly with 849 contigs and a total contig sequence length of 700.438 Mb (Table 1). Additionally, haplotype-phased contig-level assemblies were generated: a haplotype_1 assembly with 1220 contigs and a total contig sequence length of 630.117 Mb, and a haplotype_2 assembly with 1153 contigs and a total contig sequence length of 680.568 Mb. After purging haplotypic duplication from the primary contig assembly, the assembly consisted of 749 contigs and had a total contig sequence length of 671.670 Mb (Table 1). Following Hi-C-aware scaffolding, the assembly was composed of 212 scaffolds and had a total scaffold sequence length of 671.726 Mb (Table 1). Following manual curation, the assembly had 219 scaffolds, including the three chromosome-scale scaffolds expected of mosquitoes, and a total scaffold sequence length of 671.695 Mb (Figure 2A-C; Table 1). After gap-closing, the assembly had 219 scaffolds and a total scaffold sequence length of 671.687 Mb (Figure 2A,B; Table 1).

The *Ae. togoi* genome assembly is notably smaller than most other *Aedes* mosquito genomes (Figure 2D); however, we expect the genome assembly to be complete due a high BUSCO completeness score and the similar genome size estimated through k-mer analysis (Supplementary Figure 1). A scan for BUSCO single-copy orthologues estimated genome completeness to be high with low levels of duplication, with an overall BUSCO score of 97.0% (96.3% complete and single-copy, 0.7% complete and duplicated, and 0.9% present but fragmented) (Table 1).

### Synteny with other mosquito genomes

The order of orthologous BUSCO genes in *Ae. togoi* was used to infer genome synteny with other *Aedes* species. Much of the *Ae. togoi* genome is composed of large blocks that are syntenic to the *Ae. notoscriptus, Ae. aegypti*, and *Ae. albopictus* genomes, either in the same orientation (shown in blue) or inverted (shown in red) (Figure 2D). The *Ae. togoi* genome appears to match closely with the chromosome structure of *Ae. notoscriptus* (Figure 2D). The exception is at chromosome 2, where between the *Ae. togoi* and *Ae. notoscriptus* genomes there is an apparent translocation near the centromere that has caused the chromosome arms to switch sides (Figure 2D).

### Repeat sequence prediction

Repeat content was estimated to represent 63.71% of the *Ae. togoi* genome, with 17.67% as DNA transposons, 13.88% as retroelements (LTRs and LINEs), and 30.78% as unclassified repeat regions (Figure 3A; Supplementary Table 2). Repeat regions within each element type are relatively evenly distributed across the three chromosomes; however, in DNA transposons there is a noticeable decrease in abundance around centromeric and telomeric regions (Figure 3D), while the abundance of LTR (Figure 3E) and LINE (Figure 3F) retrotransposons spike around centromeric regions.

**Figure 3.**
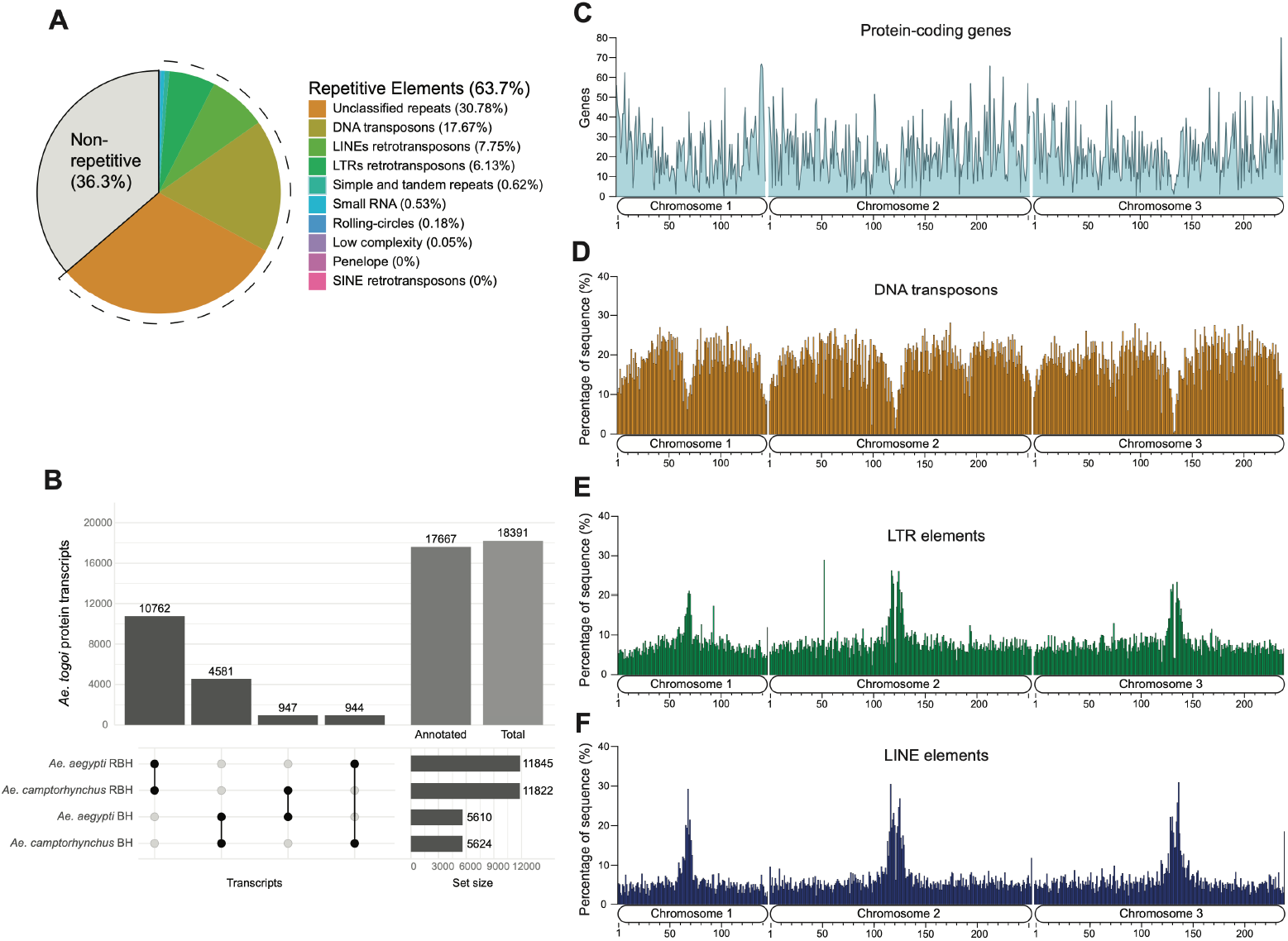
Annotation of protein-coding genes and repetitive elements in the *Aedes togoi* genome. **(A)** Repetitive element composition of the *Ae. togoi* genome, estimated using RepeatModeler and Repeat-Masker. The pie chart shows all repetitive content in the genome, separated by repetitive element classification. **(B)** Upset plot outlining the annotation of *Ae. togoi* proteins based on BLAST alignment to previously existing annotations in *Ae. aegypti* and *Ae. camptorhynchus*. Counts for predicted protein transcripts (Total), protein transcripts with an annotation (Annotated), protein transcripts matched through reciprocal best-hit (RBH) or best-hit (BH), and combinations of annotation match types are shown. Proportion and distribution of **(C)** protein-coding genes, **(D)** DNA transposons, **(E)** long terminal repeat (LTR) retrotransposons, and **(F)** long interspersed nuclear element (LINE) retrotransposons along *Ae. togoi* chromosomes. **(C-F)** Percentage of sequence values were calculated by the proportion of sequence taken up by genes and repeat elements within 1Mb stretches along the chromosomes.

### Protein-coding gene set prediction

To inform the prediction of protein-coding gene models, we generated poly-A selected RNA-seq data from adult heads, adult bodies, whole larvae, dissected larval anal papillae, and dissected larval salt gland. The BRAKER3 automated annotation tool predicted 18,391 protein-coding transcripts encoded by 12,977 genes (Table 2). 12,917 of these protein-coding genes (99.5%) were placed relatively evenly distributed on the three chromosome scaffolds, leaving 60 (0.5%) across the remaining genomic scaffolds (Figure 3C). BUSCO scores of the proteins encoded by these gene model predictions were similar to those measured at the genome scale, with 98.0% (3220 of the 3285) proteins identified as present and complete (Table 2). Of these, 3195 had single-copies and 25 were duplicated, while the remaining BUSCOs included 5 fragmented and 60 missing (Table 2).

### Functional annotation of protein coding genes

Functional annotation of the *Ae. togoi* gene set was performed by assigning GO terms, PFAM domains, and inferring orthologs based on protein sequence similarity to *Ae. aegypti* and *Ae. camptorhynchus*. This resulted in 16,819 transcripts with GO term and PFAM annotations and 17,667 transcripts with orthology annotations (Figure 3B; Supplementary Table 5). Using *Ae. aegypti* proteins as a reference database for whole-transcriptome BLAST analyses, 11,845 transcripts matched through reciprocal best hit (RBH) and were identified as putative 1:1 orthologues, while an additional 5,610 transcripts matched through best-hit (BH) as putative orthologs (Figure 3B). Using *Ae. camptorhynchus* proteins as reference, 11,822 orthologues were identified through RBH BLAST and an additional 5,624 putative orthologs were identified through BH BLAST (Figure 3B). 10,762 of these *Ae. togoi* protein annotations identified through RBH had matches to both reference species, while 221 had annotation matches only to *Ae. aegypti* and 212 had annotation matches only to *Ae. camptorhynchus (*Figure 3B).

### GO term enrichment within the specialized salt-secreting gland of the larval hindgut

Unlike other saline-tolerant mosquitoes in the sub-genus *Ochlerotatus*, whereby the salt-secreting gland is a specialized segment situated just posterior to the typical rectum, the salt-secreting gland of *Ae. togoi* larvae is an elaboration of the anal canal and is not associated with the rectum (Asakura, 1980; Asakura, 1982). (Figure 4A). To uncover the specific ion-transport mechanisms for urine concentration by the salt-secreting gland, we generated transcriptomes of this specialized organ from larvae reared in freshwater (FW) or seawater (SW). To identify transcripts enriched in the salt-secreting gland relative to the whole larvae, we also generated transcriptomes using tissues from FW- and SW-reared whole larvae. This is the first transcriptome of the specialized salt-secreting gland for a saline-tolerant mosquito larva.

**Figure 4.**
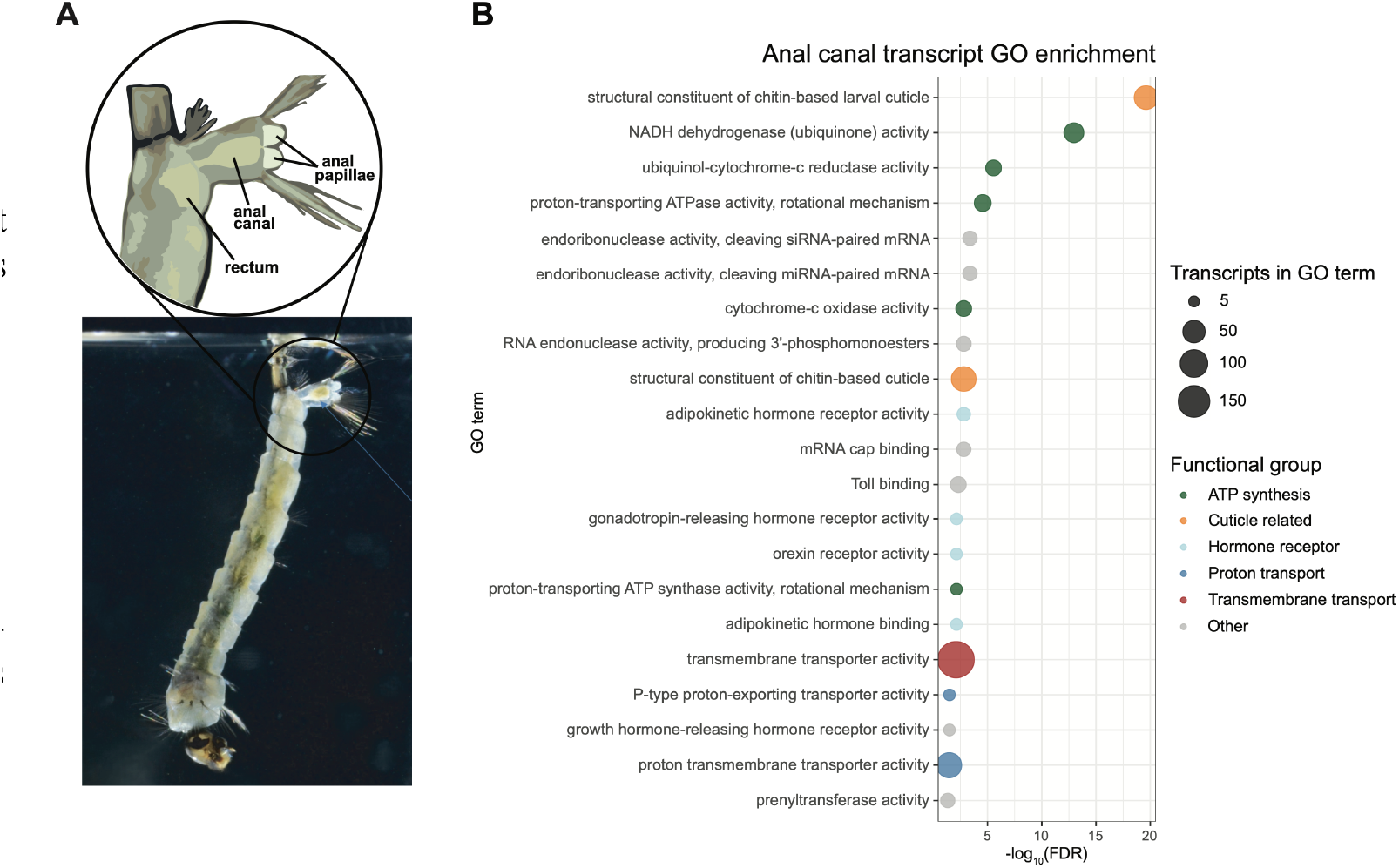
The anal canal of *Aedes togoi* acts as the primary site of salt ion excretion. **(A)** The elaborated anal canal in *Ae. togoi*, the specialized salt-gland structure, in distinction from the rectum and anal papillae ion regulatory structures. **(B)** Molecular function (MF) GO-terms enriched amongst transcripts with high expression in the anal canal (salt-gland), compared to the transcripts expressed in the anal canal and whole larvae. Significance of enrichment is measured by false discovery rate (FDR), -log_10_ transformed in the plot. The enriched GO-terms are grouped together by broad functional categories: ATP synthesis, Cuticle related, Hormone receptor, Proton transport, Transmembrane transport, and Other.

GO-term enrichment analysis for transcripts that are highly expressed in the salt gland transcriptome, compared to the whole larval transcriptome, yielded 21 significantly enriched GO-terms (Figure 4B; Supplementary Table 6A). Terms related to larval cuticle structural constituents (GO:0008010, GO:0005214), ATP synthesis (GO:0008137, GO:0008121,GO:0004129, GO:0046933, GO:0046961), proton transport (GO:0008553, GO:0015078), and transmembrane transport (GO:0022857) were identified, pointing to potential molecular functions occurring in the salt gland and its overarching role in ion regulation (Figure 4B; Supplementary Table 6A). Several GO-terms related to hormone receptor activity were also identified (adipokinetic hormone receptor activity (GO:0097003), gonadotropin-releasing hormone receptor activity (GO:0004968), orexin receptor activity (GO:0016499), adipokinetic hormone binding (GO:0097004), and growth hormone-releasing hormone receptor activity (GO:0016520)), pointing to potential hormonal regulation of salt-response mechanisms (Figure 4B; Supplementary Table 6A).

To begin to elucidate the broad impacts of salinity on gene expression in the salt gland specifically, we conducted GO-term enrichment of salt gland transcripts of larvae reared in FW and SW. Comparing SW and FW rearing conditions, we identified nine GO-terms significantly enriched in the SW condition, five of which are related to transmembrane transport activity (Supplementary Table 6C). Further, we find that only one of these GO-terms (voltagegated potassium channel activity; GO:0005249) is enriched among up-regulated transcripts in SW conditions, compared to up-regulated transcripts in the FW condition in the salt gland (Supplementary Table 6E).

### Impacts of high salinity on transcript expression in whole larvae and salt gland

Transcript expression from larvae reared in FW versus SW conditions were used to identify responses to salinity stress in the whole animal and within the salt gland, the site of hyperosmotic urine formation, specifically. At the whole larvae level, there were no significantly enriched GO-terms among transcripts in the SW condition compared to the FW condition (Supplementary Table 6B). However, there was noticeable differential expression in whole larvae reared in SW: 1014 transcripts were differentially expressed, 641 down-regulated, and 373 up-regulated (Supplementary Figure 6A; Supplementary Table 7A). The two up-regulated transcripts with the highest log2FoldChange were sodium/potassium transporting ATPase subunits (g7279.t1 and g7279.t9), both with low-moderate TPM expression in SW and zero TPM expression in FW (Supplementary Table 7A). From a sub-list of 314 transcripts relevant to ion regulation and osmoregulation, 28 were differentially expressed in whole larvae reared in SW (Table 3; Supplementary Figure 6C). Although potentially related to ion regulation in epithelia of osmoregulatory organs, this cannot be confirmed using the whole larval tissue dataset, due to the lack of tissue-specificity. Therefore, we focused our attention primarily on the salt gland transcript expression dataset for insights on ion regulatory responses to salinity stress.

Through differential expression analysis of the *Ae. togoi* salt gland in FW and SW we have identified 886 transcripts that are differentially expressed (Supplementary Figure 6B; Supplementary Table 7B). We also examined differential expression of transcripts relevant to ion regulation and osmoregulation in the salt gland, finding 11 down-regulated and 8 up-regulated (Table 3; Supplementary Figure 6D).

We further examined expression of these ion regulatory transcripts in the salt gland, in both FW and SW conditions, inclusive of ion pumps and enzymes, transporters, ion and water channels, and paracellular junctions (Figure 5A-E; Table 3).

**Figure 5.**
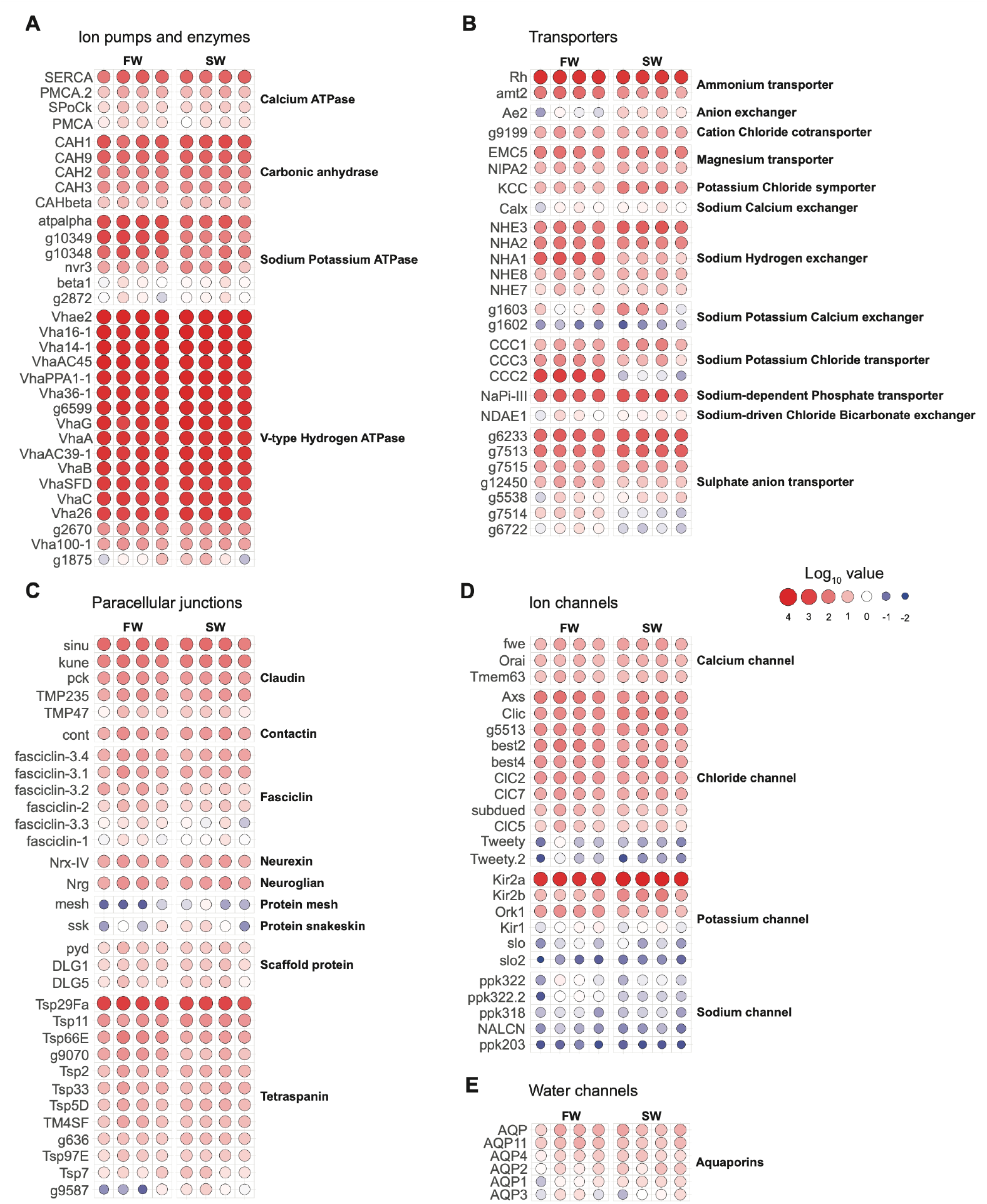
Abundance of ion regulatory-related transcripts in the salt gland of *Aedes togoi* larvae. Ion-regulatory transcripts in the salt gland of freshwater (FW) and 100% seawater (SW) reared *Ae. togoi* larvae, organized by **(A)** ion pumps and enzymes, **(B)** transporters, **(C)** paracellular junctions, **(D)** ion channels, and **(E)** water channels. **(A-E)** Each group (FW and SW) features four replicates for all genes, showing transcript abundance represented by the log TPM value (log_10_ of TPM > 0.01) scaling with both size and colour.

Among ion ATPases and other key enzymes, both calcium ATPases and carbonic anhydrases showed high expression with low differential expression across the FW and SW conditions (Figure 5A; Supplementary Table 8A). Similarly, multiple subunits of the vacuolar-type hydrogen ATPase (VHA) were very highly expressed in both FW and SW conditions. On the other hand, several subunits of sodium-potassium ATPase (NKA) (atpalpha, g10329, g10348, and nvr3) showed high expression in FW conditions and decreased expression in SW conditions, while others showed consistently low expression (beta1 and g2872) (Figure 5A; Supplementary Table 8A).

Many ion transporters show similar expression in the salt gland of larvae reared in FW and SW conditions, including the ammonium transporters, cation chloride cotransporters, magnesium transporters, sodium potassium calcium exchangers, and sodium chloride bicarbonate exchanger (Figure 5B; Supplementary Table 8B). Interesting exceptions are the potassium-chloride-symporter (KCC), sodium-hydrogen antiporters and exchangers (NHA and NHE), and sodium-potassium-chloride-cotransporters (CCC) (Supplementary Figure 6D). KCC is lowly expressed in FW but is up-regulated 2.5-fold when larvae are reared in SW. Among sodium-potassium-chloride-cotransporter paralogs, expression differs significantly; CCC1 is significantly up-regulated in SW while CCC2 and CCC3 are significantly down-regulated in SW compared to development in FW. CCC2, in particular, exhibits a significant decrease in transcript expression, from an average count of 830 TPM when in FW to 0.5 TPM when larvae are reared in SW (Figure 5B; Supplementary Table 8B). Paralogs of the sodium-hydrogen exchanger (NHA/NHE) are differentially expressed in SW, notably with NHE2 up-regulated by 1.6-fold and NHA1 down-regulated by 5-fold in SW. Additionally, both the sulphate-anion transporters (g6233 and g7513) and a phosphate transporter (NaPi-III) are highly expressed in SW and are moderately up-regulated with a 0.7 and 0.5-fold increase, respectively (Figure 5B; Supplementary Table 8B).

Cell-to-cell occluding junctions, or septate junctions, of invertebrate epithelia are a significant route for selective, transepithelial water and solute flux between cells. Septate junction proteins in the salt gland of larvae showed a wide range of expression but no differential expression between FW and SW conditions (Figure 5C; Supplementary Table 8C). Certain protein transcripts were highly expressed, for example, Tsp29Fa in the tetraspanin family and the claudin proteins sinu and kune (Figure 5C; Supplementary Table 8C).

Ion channel protein expression in the salt gland did not differ between FW and SW rearing conditions (Figure 5D; Supplementary Table 8D). Sodium channels (ppk203, ppk322, ppk322.2, ppk318, and NALCN) had a low expression, while calcium channels (fwe, Orai, Tmem63) were relatively moderately expressed overall (Figure 5D; Supplementary Table 8D). Potassium channels exhibit a range of expression patterns, with Kir1, slo, and slo2 transcripts showing low expression and Kir2a, Kir2b, and Ork1 transcripts showing relatively high expression (Figure 5D; Supplementary Table 8D). The Kir2a transcript, in particular, is highly expressed in both FW and SW. In addition, potassium (K^+^) inward rectifiers (Kir) channels Kir2a and Kir2b are differentially expressed in the salt gland, with expression of these two transcripts increasing in SW by 1.7 and 1.5-fold, respectively. Specific chloride channels are also moderately expressed in the salt gland, except Tweety and Tweety-like which have a low transcript expression (Figure 5D; Supplementary Table 8D).

Water channels (i.e. aquaporins) were not differentially expressed in the salt gland between FW and SW conditions and none of the aquaporin transcripts were highly enriched in the salt gland (Figure 5E; Supplementary Table 8E).

## Discussion

Here, we present a *de novo* reference genome assembly for *Aedes togoi*, the first chromosome-scale assembly for a saline-tolerant mosquito. In addition, we provide transcriptomic insights into the mechanism by which this species tolerates high salinity and provide a putative model of salt secretion in the novel anal canal salt gland. With these, we hope to support further studies that expand our understanding of mosquito genome evolution, develop genetic toolkits in lesser-studied branches of the *Aedes* genus, and characterize saline-tolerance strategies and mechanisms in primarily freshwater organisms.

### Unexpected chromosome arrangement

Advances in genome sequencing technology, and heightened efforts to sequence a broader range of species, have transformed the way we can investigate mosquito genome organization. We used synteny to related *Aedes* species and the ability of Hi-C data to identify points of mis-assembly to validate chromosome structures following scaffolding. Following scaffolding and manual curation, the chromosomal structure of our *Ae. togoi* genome assembly matches the overall structure of the *Ae. notoscriptus* (GCA_040801935.1) and *Ae. triseriatus* (personal communication with John Soghigian) genomes, based on BUSCO gene-order showing syntenic arrangement of large blocks between these *Aedes* species (Figure 2D). The three largest scaffolds HiC_scaffold_101, HiC_scaffold_100, and HiC_scaffold_109, are observed to be chromosomes 1, 2, and 3, respectively (naming matched to *Ae. notoscriptus* chromosome numbers) (Figure 2D). There is one structural rearrangement in HiC_scaffold_109 that is not reflected in the matching *Ae. notoscriptus* chromosome 2, but it instead aligns with the arrangement of the *Ae. triseriatus* chromosome 2 (personal communication with John Soghigian) and is supported by our *Ae. togoi* Hi-C contact map signal patterning (Figure 2C; Supplementary Figure 4E). The concentration of alignment signal within scaffolds along the diagonal, other than the small regions of alignment signal between repetitive regions believed to be the chromosomal centromeres and telomeres (Figure 2C), and the lack of off-diagonal alignment signal, supports the scaffolding and manual rearrangement of chromosomes within our *Ae. togoi* genome assembly (Figure 2C; Supplementary Figure 4).

Chromosomal rearrangements similar to the *Ae. togoi/Ae. notoscriptus* chromosome 2 arm translocation are not uncommon within the *Aedes* genus and mosquitoes as a whole (Morinaga et al., 2025; Ryazansky et al., 2024a). For instance, comparing the previously assembled *Ae. aegypti* and *Ae. notoscriptus* genomes indicates large-scale rearrangement of chromosome arms (Figure 2D). Notably, syntenic blocks on the arms of chromosomes 2 and 3 are swapped with each other between these species, with half of *Ae. aegypti* chromosome 2 seemingly matching to *Ae. notoscriptus* chromosome 2, and the other half matching to *Ae. notoscriptus* chromosome 3 (Figure 2D). Additional highquality genome assemblies from more *Aedes* species are required to have a better sense of how chromosomal rearrangement has played a role in the radiation of the *Aedes* genus.

### Comprehensive gene-set annotations

A high BUSCO score of 98.0% for the predicted proteome sequences gives us confidence that the large majority of encoded *Ae. togoi* proteins are captured by our gene-set predictions. Further, with 96.1% (17,667 out of 18,391) of predicted protein transcripts matching to previously annotated proteins in neighbouring species (Figure 3B; Table 2), we gain a baseline, sequence-level understanding of proteome function in this emerging study organism. Because there are so few saline-tolerant mosquitoes, and so few of these have high-quality genomic and transcriptomic resources, these annotations represent key points of reference into the landscape of saline-tolerance evolution in mosquitoes, potentially highlighting genomic similarities and protein characteristics common between saline-tolerant species. Therefore, these annotations provide a useful resource for future analyses characterizing specific genes and proteins and tracing the evolution of gene families, within the lens of understanding saline-tolerance lineages in particular.

### Relatively small genome size driven by reduced repeat content

At just 0.67 Gb, the *Ae. togoi* genome is among the smallest *Aedes* genome assembled so far (Supplementary Table 4). Compared to *Ae. aegypti*, which we estimated to have roughly 75.30% of its genome composed of repetitive content, the 63.71% repetitive content of the *Ae. togoi* genome represents a roughly 534.9 Mb (534,962,813 bp) reduction in genomic repeat sequence (Supplementary Figure 5; Supplementary Table 2). Therefore, the genomic sequence length discrepancy between these two species (roughly 607 Mb) largely seems to be due to reduced repetitive content in *Ae. togoi*. Genome size is known to vary widely between species, and even within species between sub-populations, often due to differences in repetitive content (Black and Rai, 1988; McLain et al., 1987). The relatively large genomes of *Aedes* mosquitoes are a prime example of this as it is believed the ballooning of repetitive elements has caused *Aedine* species like *Ae. aegypti* and *Ae. albopictus* to have much larger genomes than species in other mosquito families like *Anopheles* and *Culex* (Habtewold et al., 2024; Matthews et al., 2018; Palatini et al., 2020; Ryazansky et al., 2024b). This genome assembly for *Ae. togoi*, as a smaller *Aedes* genome with relatively low repetitive sequence content, adds an important reference point for understanding the dynamics of repetitive elements and their effects on fitness traits like transcriptional gene regulation and phenotypic adaptability.

Transposable elements (TEs) in particular have been linked to genome size variation, genome rearrangements, and phenotypic adaptability to new environmental conditions, although their role of these in mosquitoes has only recently started to be outlined (Daron et al., 2025; Schrader and Schmitz, 2019). Within Culicine mosquitoes, TE abundance is known to be high, with trends of retrotransposons being predominant over DNA transposons and there being an even distribution of TEs along chromosomes (Daron et al., 2025; Melo and Wallau, 2020). Alternatively, we see a that *Ae. togoi* DNA transposons (17.67% of genome) make up a greater proportion of genomic sequence than retrotransposons (13.88% of genome) (Figure 3A), in severe contrast *Ae. aegypti* which (based on estimates with our described methods) has a strong leaning towards retrotransposons (44.57% of genome) over DNA transposons (1.3% of genome) (Supplementary Figure 5). There is also a distinction in the distribution of retrotransposons (LINEs and LTRs) and DNA transposons in *Ae. togoi*, in which retrotransposons have greater abundance at centromeric regions (Figure 3E,F) while DNA transposons have reduced abundance at centromeric and telomeric regions (Figure 3D). These differences in TE abundance and distribution may signify differences in the evolutionary history of *Ae. togoi* compared to other *Aedes* species studied so far.

### Transposable elements and chromosomal inversions in adaptive trait evolution

Harsh environmental conditions may have left their mark on *Ae. togoi* genome as the species developed towards their coastal rockpool niche. TE activity has been implicated as a mechanism for rapid adaptation required to explain the genetic paradox of invasive species, in which species are able to rapidly adapt to new environments despite genetic bottlenecks (Schrader and Schmitz, 2019; Stapley et al., 2015). Further, it has been shown that stressful environmental conditions can trigger TE activity (Horváth et al., 2017). TEs have previously been directly linked to the emergence of adaptive phenotypic traits such as insecticide resistance in *An. funestus* and cold-weather tolerance in *D. montana* (Mugenzi et al., 2024; Tahami et al., 2024). These together point to the potential that cold-weather and fluctuating water temperature and salinity in temperate, coastal regions may have played a similar role in triggering TE activity and rapid adaptation in *Ae. togoi*.

Chromosomal inversions have also been identified as influential in the process of adaptive trait evolution and even speciation (Berdan et al., 2022; Feder and Nosil, 2009; Hoffmann et al., 2004; Jay and Joron, 2022; Sharakhova and Sharakhov, 2025; Stump et al., 2007) Inversions support adaptation to new environmental conditions and provide fitness advantages to stressors by effectively reducing recombination rate within the inverted region and increasing the linkage between potentially beneficial alleles (Berdan et al., 2023; Jay et al., 2018). In mosquitoes, inversions have been linked to specific adaptive phenotypic traits like drought tolerance, insecticide resistance, anthropophilic feeding behaviour, and changes to larval habitat (Ayala et al., 2014; Ayala et al., 2019; Fontaine et al., 2015; Ingham et al., 2021; Love et al., 2019; Main et al., 2016; Sharakhova and Sharakhov, 2025; Small et al., 2023; Weetman et al., 2018). What’s more, it seems that specific environmental conditions can drive distinct patterns of chromosomal inversions, resulting in the selection of similar traits affecting fitness in that environment (Liang et al., 2025; Ma et al., 2024). It’s possible that chromosomal inversions have played an important role in the speciation and adaptation of *Ae. togoi* to tolerate cold weather and high salinity.

Further, there are potential interactions between TEs and chromosomal inversions, with evidence that TE activity promotes the occurrence of chromosomal inversions and possibly other types of chromosomal rearrangements (Kent et al., 2017). With the abundance of TEs and chromosomal inversions in mosquito genomes it may be that these are primary and interactive drivers of rapid adaptation in *Ae. togoi* and other mosquitoes. This chromosome-level genome assembly provides a resource to study these possible interactions, although further work comparing chromosome structure and repetitive element composition of many more mosquito species are required to fully understand this dynamic.

### Molecular investigation of the *Ae. togoi* salt gland for active salt secretion and urine concentration

Numerous elegant and thorough studies over the past 80 years have implicated the salt gland of euryhaline mosquito larvae as the primary site of active salt secretion from the blood and into the urine to void excess salt as a strategy to hydrate (Asakura, 1980, Patrick et al., 2024; Bradley and Phillips, 1975; Bradley and Phillips, 1977). Here, we provide mechanistic support for these findings with transcriptomic evidence and present a putative molecular model for ion flux across the salt gland epithelium.

GO-terms enriched in the salt gland transcripts highlight molecular functions important to the overarching role of this critical organ in salt excretion. Enriched GO-terms were centered around larval cuticle composition, ATP synthesis, proton transport, and transmembrane transport, which together provide clues into how *Ae. togoi*’s extreme salt tolerance may be achieved (Figure 4B; Supplementary Table 6A). It is well established that integumental adjustments of saline-tolerant insects include reduced permeability of the cuticle to attenuate water loss (Nicolson and Leader, 1974), and transcripts related to larval cuticle structure in our datasets highlight the importance of the cuticle as a barrier to avoid desiccation (Wigglesworth, 1933). Similarly, the enrichment of transmembrane transport transcripts was unsurprising and demonstrates the presence of ion transport machinery necessary to drive active salt secretion via the salt gland when larvae develop in seawater. Enrichment of proton transport GO-terms are indeed exciting and suggest parallels with known mechanisms of hyperosmotic urine formation of *Ae. taeniorhynchus* larvae via H^+^-driven sodium secretion in the salt-secreting portion of the rectum (Patrick et al., 2024). The high transcript expression of mitochondrial ATP-synthase subunit transcripts and V-type proton ATPase subunit transcripts (Supplementary Table 6A) together that point towards proton gradients generated by proton ATPases as a mechanism for powering ion transport.

Transcripts of several hormone-related GO-terms are highly enriched in the salt gland; adipokinetic hormone (AKH) receptor activity, gonadotropin-releasing hormone receptor activity, orexin receptor activity, adipokinetic hormone binding, and growth hormone-releasing hormone receptor activity. The enrichment of these GO-terms comes almost entirely from enrichment of g10702 transcripts g10702.t1-g10702.t5 (Supplementary Table 6A). Interestingly, these transcripts are annotated as AKH/corazonin-related peptide (ACP) receptor transcripts when referenced to *Ae. camptorhynchus* proteins but are annotated as gonadotropin-releasing hormone II receptor transcripts when referenced to *Ae. aegypti* proteins (Supplementary Table 5). The ACP receptor is a structural intermediate between AKH and corazonin, which are two distinct insect neuropeptides (Afifi et al., 2023; Hansen et al., 2010). AKH is largely involved in energy homeostasis (Gade et al., 1997) while corazonin has been shown to have a dual role in regulating metabolic and osmotic stress (Kubrak et al., 2016; Zandawala et al., 2021). ACP is localized to the brain of adult *Ae. aegypti* with axonal projections to the abdominal ganglia which are enriched in ACP receptor transcripts (Afifi et al., 2023; Wahedi and Paluzzi, 2018). This is suggestive of a potential role of ACP in the regulation of release of other neuropeptides that are secreted from neurosecretory cells of the abdominal ganglia, specifically those with important roles in diuretic or anti-diuretic signalling pathways (Chen et al., 1994; Sajadi et al., 2020). The enrichment of a putative ACP receptor in salt gland tissue of larvae warrants a deeper investigation and might signify an important regulatory pathway when larvae are required to transition between antidiuresis and diuresis states as external salinity fluctuates.

Differential expression analyses of salt gland transcripts from larvae reared in FW and SW conditions, aiming to identify candidate ion transport processes critical for hyperosmotic urine production during high salinity, provided little information (Supplementary Table 7). In fact, many transcripts that are implicated in active ion secretion in renal and extra-renal ion-transporting epithelia of insects are enriched in the salt gland of *Ae. togoi* larvae regardless of external salinity. Nonetheless, raw transcript expression (TPM) of genes revealed that many ion channels, transporters, and pumps are always highly enriched in this specialized organ. For example, V-type proton ATPase (VHA) subunits are very highly expressed in both FW and SW conditions (Table 3). This pump provides the driving force for transepithelial Na^+^ secretion from the hemolymph into the salt gland lumen of euryhaline *Ae. taeniorhynchus* larvae and these data strongly suggest a parallel mechanism for Na^+^ secretion via the salt gland of *Ae. togoi* (Patrick et al 2024). Of course, transcript expression does not necessarily represent changes in VHA pump activity and transcriptional regulation may not be the primary method for responding to changes in ion and osmo-regulatory needs (Patrick et al., 2024). In the renal Malpighian tubules of the *Aedes aegypti* mosquito, VHA protein complexes are dissociated to achieve antidiuresis and this phenomenon is under neuroendocrine control (Sajadi et al., 2023). As such, further studies that measure specific enzyme activities and that localize cytosolic and membrane protein complexes spanning dilute and hypersaline salinities, are required. Post-translational methods of regulation, for example, through reversible dissociation or phosphorylation causing protein inactivation/reactivation (Kandel et al., 2022; Kengne et al., 2019) may be productive avenues to be explored.

### Putative cellular model of salt secretion across the salt gland epithelium

Using the expression of ion-regulatory transcripts enriched in the salt gland of larvae in SW, we propose a working cellular model of transepithelial ion secretion across the salt gland epithelium of *Ae. togoi* (Figure 6). Here, we lean on the well-characterized transport models of ion flux from other mosquito osmo- and ion-regulatory organs including the Malpighian tubules, rectum, and anal papillae (Beyenbach et al., 2010; Duca et al., 2011; Piermarini et al., 2011; Piermarini et al., 2015; Piermarini, 2016; Chasiotis et al., 2016; Jonusaite et al., 2016; Durant et al., 2021).

**Figure 6.**
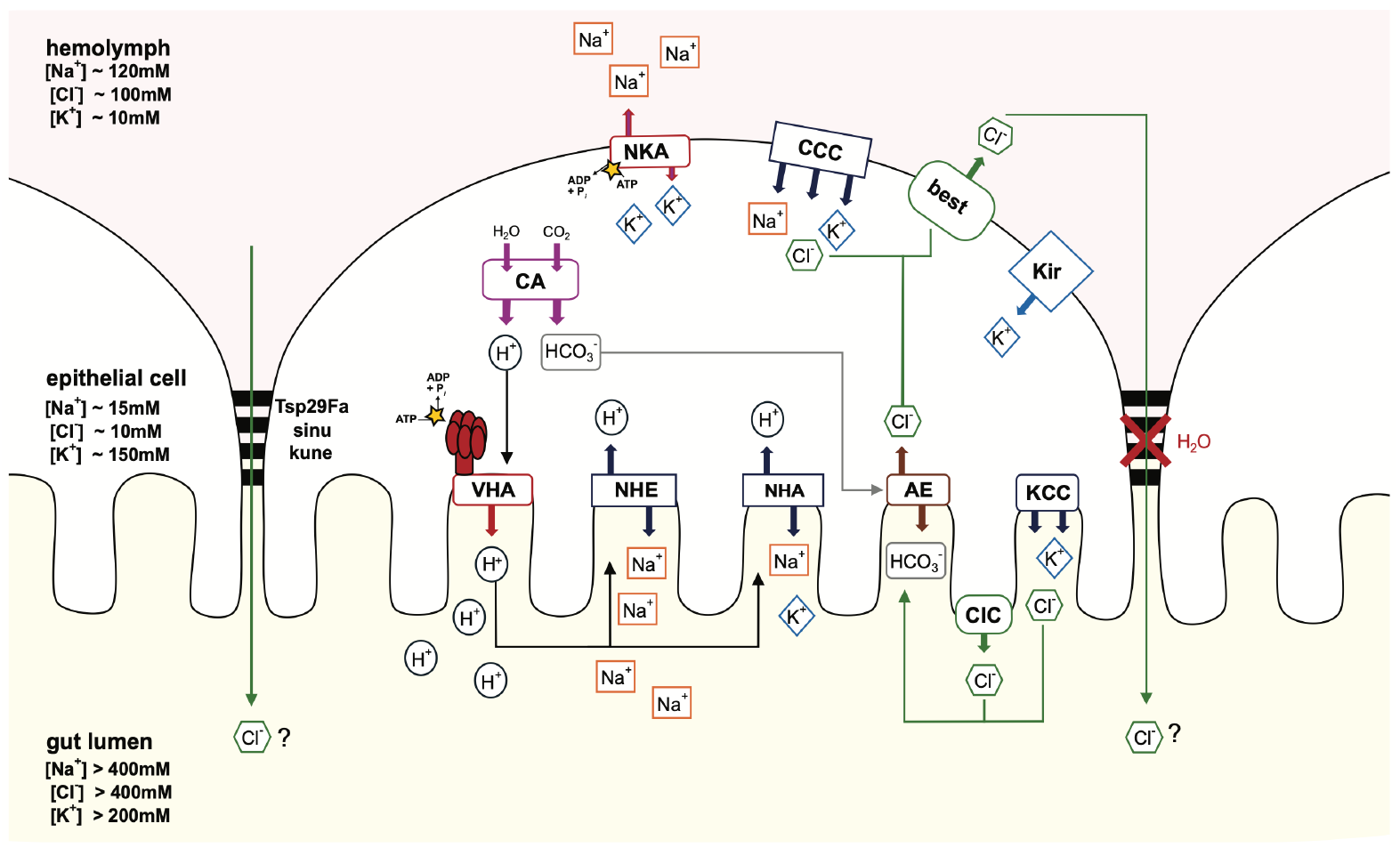
Putative model of sodium and chloride secretion in the salt gland of *Aedes togoi* larvae. Working cellular model of sodium (Na_+_) and chloride (Cl_-_) secretion by the salt gland of the hindgut of *Aedes togoi* larvae. A putative model of salt secretion from the hemolymph, across the salt-gland epithelium, and into the lumen is presented using enriched sequences. On the apical membrane, the V-type H_+_-ATPase (VHA, multiple subunits) pumps protons (H_+_) generated by carbonic anhydrases (CA, multiple isoforms) into the luminal space. High luminal H_+_ drives the secretion of mostly Na+, and some K_+_, apically into the lumen via the sodium-hydrogen exchanger (NHE) and an inwardly rectifying potassium channel (Kir) on the basolateral membrane. The potassium chloride co-transporter (KCC) and chloride channel (ClC) are low resistance routes for apical K_+_ and Cl_-_ flux into the lumen, and this is driven by the K_+_ and Cl_-_ electrochemical gradients. Cl-flux back into the cell cytosol across the apical membrane might also drive bicarbonate (HCO_3-_) secretion into the salt gland lumen transported by an anion exchanger (AE). We suggest that basolateral Cl_-_ flux back into the hemolymph from the cytosol occurs via a low resistance pathway that involves a chloride channel (best). We propose that a major route for Cl_-_ flux into the lumen might be a low resistance paracellular pathway, similar to other insect secretory epithelia. The positive luminal space created by the excess sodium ions drives the paracellular movement of chloride through water-impermeable septate junction protein complexes (sinu, kune, and Tsp29Fa). The active secretion of high levels of Na_+_, K_+_, and Cl_-_ across the salt gland epithelium, one that has a low aquaporin expression and a high resistance to water flux, allows for the generation of a hyperosmotic urine by larvae to achieve ion and water balance when in seawater.

Drawing from the mechanism for Na^+^ secretion described for the salt gland of *Ae. taeniorhynchus* (Patrick et al., 2024), we propose that vacuolar-type hydrogen pump (VHA) is expressed on the apical membrane and pumps protons towards the lumen, thereby contributing to a negative cytosolic voltage and establishing a H^+^ gradient as a driving force for the secretion of other ions (Figure 6). Given the high expression of ammonium transporters (amt2, Rh), apical expression of VHA likely facilitates an acid trapping mechanism for nitrogenous waste (NH_3_/NH_4_ ^+^) excretion from the hemolymph, as is the case for many excretory organs of mosquito larvae including the anal papillae of freshwater-restricted species *Ae. aegypti* (Chasiotis et al., 2016). We propose the protons for VHA are supplied by cytosolic carbonic anhydrase (CA) (Figure 6), given the high expression of multiple CA isoforms and previous work in *Ae. aegypti* highlighting a similar role in the anal papillae (Chasiotis et al., 2016; Duca et al., 2011; Durant et al., 2021).

While sodium-potassium-ATPase (NKA) (Figure 6) transcripts (Atpalpha) are highly up-regulated with SW rearing in whole larvae (Figure 5), NKA expression and pump activity is critical for many biological processes and its function is ubiquitous in nervous, muscle, and epithelial tissues. We include NKA in our model of salt-secretion as transcript levels of multiple subunits are enriched, but we acknowledge that the salt gland in our dataset is comprised of epithelial, muscle, and probably nervous tissue that would each have NKA transcript expression. Given the notable lack of NKA protein expression in the salt gland epithelium of *Ae. taeniorhynchus* larvae (Patrick et al., 2024), we do not rule out the possibility that NKA protein is not a critical energizer of transepithelial ion flux in the salt gland of *Ae. togoi* larvae. Instead, we propose protons drive the secretion of mostly sodium and some potassium through highly expressed apical sodium hydrogen exchangers, NHE and NHA (Figures 5 and 6), similarly to *Ae. taeniorhynchus* (Patrick et al., 2024). NHA is expressed in the stellate cells of the Malpighian tubules, suggesting NHA-mediated transport of both Na^+^ and K^+^ into the tubule lumen (Rheault et al., 2007; Beyenbach et al., 2010).

In the salt gland of *Ae. taeniorhynchus*, it was suggested that a basally expressed cation-chloride cotransporter (CCC) transports ions into the cytosol from the hemolymph (Patrick et al., 2024), similarly to the principal cells of the Malpighian tubules of *Ae. aegypti* (Beyenbach and Piermarini, 2011). Given that the different CCC isoforms have distinct expression patterns in the salt gland of *Ae. togoi* (Figure 5), it is very likely that a basally expressed CCC (CCC1) is a low resistance route for electroneutral ion flux into the cytosol from the hemolymph (Figure 6). CCC1 has been suggested to facilitate ion secretion, whereas CCC2 and CCC3 might have critical functions in absorptive epithelia (Piermarini et al., 2017). The very high expression of the inward rectifier potassium channels (Kir2a and Kir2b) (Figure 5) likely also mediate potassium flux into cytosol from the hemolymph, for secretion across the apical membrane via NHA and a potassium chloride cotransporter (KCC) (Figure 6). Kir2b is highly expressed in the principal cells of the Malpighian tubules and is critical for transepithelial potassium secretion (Piermarini et al., 2015). We propose the enriched KCC is apically expressed similarly to the renal tubules of *Ae. aegypti* for the generation of primary urine using potassium as a major osmolyte (Piermarini et al., 2011).

In this putative model of salt-secretion, apical KCC is a low resistance route for the removal of intracellular potassium towards the lumen, and also a path for chloride flux for secretion in the urine. We posit that luminal Cl^-^ is a primary driving force for HCO_3_ ^-^ secretion via an apically expressed anion exchanger (AE) (Figure 6). Intracellular bicarbonate is generated by cytosolic carbonic anhydrase (CA) and is concentrated in the cytosol as a by-product of cytosolic CA activity and the secretion of H^+^ by VHA (Figure 6). Bicarbonate ions are also concentrated in seawater that the larvae drink, and therefore, would likely need to be secreted from the hemolymph into the urine. The AE provides a low resistance route for HCO_3_^-^ secretion and this occurs via electroneutral exchange with luminal Cl-that is transported apically via the KCC and an apical chloride channel (ClC). In the anal papillae of freshwater *Ae. aegypti*, an AE is apically expressed in the epithelium along with a basal chloride channel for chloride absorption (Duca et al., 2011). Here, we suggest a similar mechanism for transcellular chloride flux from the lumen and back into the hemolymph via a basally expressed chloride channel (best). We propose that the main route of chloride secretion into the urine for excretion is paracellular via septate junctions and driven by an electrical gradient established by high luminal cation levels (Figure 6). Paracellular chloride flux has been demonstrated in the Malpighian tubules, albeit to drive fluid secretion (Beyenbach et al., 2010). Given the low expression of water channels (Figure 5) and the increased desiccation risk in seawater, we suggested that there is a very high resistance of the salt gland epithelium to transcellular or paracellular water flux (Figure 6). We also note that the septate junctional complex proteins claudins, sinu, kune, and tetraspanin (Tsp29Fa) that are enriched in the salt gland (Figure 5) play a large role in the development, regulation, and maintenance of the septa, and so they are very likely important in regulating paracellular chloride secretion (Jonusaite and Himmerkus, 2024).

## Conclusion

With the addition of the *Aedes togoi* genome assembly, accompanying protein-coding annotations, and transcriptomic datasets, we support future comparative genomics and physiological studies seeking to understand the genetic and molecular mechanisms of mosquito physiology. It has been known for almost a century that salt-secreting glands of saline-tolerant *Aedine* mosquito larvae are necessary for hydrating in seawater, but mechanistic information underlying urine concentration remains scarce. The salt gland transcriptomes and putative model of salt-secretion open new doors to a comprehensive understanding of salt-secretion that is fundamentally different to other insects in desiccating environments. These resources and findings establish *Aedes togoi* as a study species well-suited for characterizing ion regulation and salinity tolerance in mosquitoes.

## Author contributions

JC contributed to: conceptualization (genomic sequencing, genome assembly, genome annotation, transcriptomic analysis), data curation (processing, cleaning, and annotating of genomic and transcriptomic sequencing data), formal analysis (genome synteny, repeat content, gene annotation, protein orthology, transcript differential expression), investigation (genomic sequence data collection, genome assembly, genome annotation, genome content analysis, genome synteny analysis, repeat content analysis, transcriptomic analysis), methodology (genomic sequencing, genome assembly, genome annotation, genome content analysis, genome synteny analysis, repeat content analysis transcriptomic analysis), project administration, software, validation, visualization, and writing (original draft and review/editing).

EK contributed to: investigation (performed experiments and data collection related to the salt-gland transcriptomics), methodology (salt gland transcriptomic analysis and salt-secretion model) data curation (processing, cleaning, annotating of salt gland RNAseq data), visualization (preparation and creation of figures for salt gland RNAseq data and salt-secretion model) and writing (original draft for the salt gland transcriptomic analysis and salt-secretion model).

OP contributed to: investigation, methodology, and formal analysis of rock pool salinity and temperature findings.

AKPN contributed to: investigation (whole larvae RNA-seq data collection).

DAHP contributed to: methodology and supervision of rock pool salinity and temperature findings.

ACD contributed to: conceptualization (salt gland transcriptomics), funding acquisition (salt gland RNAseq), methodology (salt-secretion model), and writing (review/ editing).

BJM contributed to: conceptualization (genomic sequencing, genome assembly, genome annotation, transcriptomic analysis), funding acquisition, project administration, resources (mosquito sample collection, instrumentation, genomic sequencing, computing resources), supervision, visualization (created species distribution and rock pool salinity and temperature figures), and writing (review/editing).

## Data availability

Assembly and sequencing reads accessible on NCBI under BioProject accession PRJNA1370975 and assembly, annotation, and additional information is available at https://github.com/bnmtthws/2026_Aedes_togoi_genome

## Acknowledgements

We would like to acknowledge members of the Matthews lab who provided feedback and support throughout the course of this project, particularly Leisl Brewster and Elva Vidya who provided helpful comments on earlier versions of the text and figures. We would like to thank Nicholas K. Tochor for visualizations in figures presenting salt gland structure and the overview of genomic and transcriptomic sequencing and analysis. We would like to acknowledge technical assistance and guidance from Marco Todesco during the generation and interpretation of Hi-C data. Finally, we would like to thank John Soghigian for his open discussions of unpublished work and for his suggestion to curate *Ae. togoi* chromosome synteny based on *Ae. notoscriptus* and *Ae. triseriatus* chromosome structures.

## Materials and Methods

### Mosquito collections and rearing

All field work for this project took place in Lighthouse Park in the city of West Vancouver (49.3316° N, 123.2636° W). To investigate the factors that affect larval survival a variety of abiotic and biotic factors were investigated over the course of a year (April 2021-April 2022). Sixteen rock pools in the southwest region of the park were chosen as sample sites. These rockpools represented various potential breeding habitats for *Ae. togoi* as they varied in size, volume, salinity, temperature, and distance from the sea. All rock pools were visited once a week for one year and the salinity, pH, temperature, approximate volume, and larval abundance was measured during each visit.

The temperature and salinity of the individual rock pools were measured using a Fisherbrand salinity meter pen (catalog #15078202). pH was measured using a Fisherbrand pH pen (catalog # S35927). The technique for measuring volume of the pools was adapted from a previously published protocol (Brenha-Nunes et al., 2016). The total area of each rock pool was divided into smaller shapes (i.e, square, rectangle, triangle) and the area of these shapes was measured using a Mileseey brand laser measuring device. Two to three depth measurements were taken for each shape using a 2m fold up ruler and an average depth was calculated. The volume of each shape was calculated, and these volumes were summed to get an approximate volume for the rock pool. Because the volume of the rock pools was extremely variable, volume was recalculated each week. Depths were taken from the same point each week to ensure that how the volume was changing week to week was being accurately measured. Historic weather data such as temperature and precipitation were obtained from Environment and Climate Change Canada Meteorological Service of Canada (ECCC-MSC) at the Vancouver Harbour CS weather station (CWHC). The total precipitation in the previous week, previous two weeks, and previous three weeks were calculated and used as variables. Similarly, the temperature from the previous week, previous two weeks, and previous three weeks were summed to create historic temperature variables. Finally, a mosquito dipper (volume 300ml) was used to measure relative larval and pupal abundance. The pool was visually inspected to see where the largest density of larvae and pupae was, and this is where the measurement was taken. A total of three dips were taken; larvae and pupae were replaced after each dip and the average number of larvae and pupae per dip was calculated.

All *Ae. togoi* mosquitoes used in this study were retrieved from our lab colony, originally collected from supra-littoral rock pools in Lighthouse Park in West Vancouver (Figure 1A) with permission granted from the District of West Vancouver.

### High Molecular Weight (HMW) Genomic DNA extraction, library preparation, and sequencing

To generate HiFi reads, HMW DNA was extracted from a single male *Ae. togoi* pupa using a modified version of the NEB Protocol for High Molecular Weight (HMW) DNA Extraction from Tissue, utilizing changes to reagent volumes outlined in the ‘Using Very Low Tissue Amounts’ section of the kit (T3060) product manual. The pupa was snap-frozen in dry ice and then gently ground using only up and down strokes, using a micro-pestle. 50µL of NEB Tissue Lysis Buffer was added while grinding continued to break up any large pieces. Another 50µL of NEB Tissue Lysis buffer was added to rinse tissue on the pestle into the tube. 10µL of NEB proteinase K (800U/mL) and 4µL of 10% SDS solution were added before incubating the sample at 56°C for one hour. The thermomixer used for incubation was set to 500rpm agitation for the first 15 minutes, while the remaining 45 minutes had no agitation. 5µL of NEB RNase A solution (20mg/mL) was added before incubating the sample again at 56°C for 10 minutes with 300rpm agitation. 55µL of NEB Protein Separation Solution was added before mixing the sample by slow repeated inversion of the tube for one minute. The sample was centrifuged at 16,000 x g for 30 minutes, at 20°C. The clear, upper fraction of the sample containing DNA was transferred to another tube using a wide-bore pipette tip. Two NEB DNA Capture Beads and 90µL of isopropanol were added to the sample before mixing on a vertical rotating mixer at 10rpm for five minutes. Our protocol then continued following the steps listed in the Monarch High Molecular Weight DNA Extraction Kit for Tissue manual, starting from Step 3 of ‘Part 2 HMW gDNA Binding and Elution’ section. Library preparation using the ultra-low DNA input workflow and PacBio HiFi sequencing on two SMRT PacBio Sequel II flow-cells were carried out by the University of Oregon’s Genomics and Cell Characterization Core Facility (GC3F). Reads from this dataset can be found at NCBI SRA (Supplementary Table 1).

Nanopore reads were generated from four libraries, prepared from four *Ae. togoi* samples: a single female pupa, a single male pupa, a pool of 8 female pupae, and a pool of 8 male pupae. HMW DNA were extracted from each sample using the same protocol as outlined above for HMW DNA extraction for HiFi sequencing, except reagent volumes used differed for the pooled samples, on account of additional tissue in the sample. The reagent volumes used for the two pooled samples were: 300µL of NEB Tissue Lysis Buffer total (150µL while grinding, 150µL for rinsing), 5µL of 10% SDS solution, 10µL of NEB Proteinase K solution, 5µL of NEB RNase A solution, 150µL of NEB Protein Separation Solution, 275µL of isopropanol, and 62µL of elution buffer (10mM TRIS-HCL, 0.1mM EDTA; pH 8.0). ONT library preparation and sequencing was carried out by the BC Genome Science Centre, following standard Nanopore PCR-free multiplexed genome library construction without shearing, using one PromethION flow cell. Reads from this dataset can be found at NCBI SRA (Supplementary Table 1).

### Hi-C genomic DNA extraction and library preparation

Hi-C data was collected from a single male *Ae. togoi* pupa, following a modified protocol adapted from the Arima Genomics Arima High Coverage HiC Kit for Animal Tissues (Material Part: A410110; Document Part: A160162 v01) and Arima Library Preparation Module for Arima High Coverage Kit (Material Part: A101030, A303011; Document Part: A160186 v02) manuals. The sample was snap-frozen using liquid nitrogen, pulverized into a fine powder using a micro-pestle, and then mixed with 1mL of 1X PBS solution. Our protocol continued following the steps listed in the Arima High Coverage HiC Kit manual starting from step 2 of ‘Crosslinking – Low Input’, continuing through to the HiC, quality control, and library preparation sections as stated. Cross-linked DNA was fragmented to the target 550-600bp range using a ME220 Covaris Focused-ultrasonicator with the following parameters: 40 watts power, 10% duty factor, 1000 cycles per burst, 20 °C water temperature, 75 second duration. The Hi-C library was sequenced using one P2 flow cell on the Illumina NextSeq2000, aiming for 150x-depth, pairedend, 150bp length reads. Reads from this dataset can be found at NCBI SRA (Supplementary Table 1).

### RNA extraction, library preparation, and sequencing: adults

mRNA from adult *Ae. togoi* males and females were profiled by separately collecting heads and bodies from four males and five females and preparing for RNA extraction by adding 450µL of TRIzol LS and 150µL of PBS solution to each sample of pooled mosquito heads or bodies. Samples were disrupted using an Omni Bead Ruptor Elite 19-040E at 6.0m/s for 30 seconds and then incubated at room temperature for 5-10 minutes. 130µL of chloroform was added before homogenizing by vortex at max speed for 20-30 seconds and allowing to incubate at room temperature for 10 minutes for phase separation to occur. Samples were centrifuged at 15,000 *x g* for 15 minutes at 4°C. The aqueous phase of each sample was transferred to new tubes and then ∼250µL of isopropanol was added. Samples were left to incubate at –20°C overnight. The next day, samples were centrifuged at 10,000 *x g* for 10 min at 4°C. Supernatant was removed from the tube, leaving the pellet behind. 1mL of ice-cold ethanol was used to wash the pellet. Samples were mixed by vortex briefly. Samples were then centrifuged at 7,500 x g for 5 min at 4°C. Ethanol was removed from the tubes and then the tubes were left to air-dry for 3-5 minutes at room temperature. The RNA pellet was dissolved by adding nuclease-free water and then letting the samples sit on ice for 1 hour. All samples were the mixed by vortex briefly before storing at -80°C. RNA samples were sent to Novogene for library preparation and sequencing. Each sample went through poly(A) enrichment and standard library preparation protocols. The libraries were sequenced on the Illumina NovaSeq X Plus series for paired-end, 150bp length reads. Reads from this dataset can be found at NCBI SRA (Supplementary Table 1).

### RNA extraction, library preparation, and sequencing: whole larvae

mRNA was collected from whole *Ae. togoi* larvae reared in freshwater and saltwater conditions. Briefly, larvae were hatched and then reared in either dechlorinated Vancouver tap water or dechlorinated Vancouver tap water supplemented with increasing concentrations of artificially formulated seawater (Instant-Ocean; increasing by the equivalent of ∼100% sea-water per day, final concentration: ∼300% seawater), until they reached 4^th^ instar. Larval samples were dried and then placed in 350µL of Qiagen RNeasy Lysis Buffer and 3.5µL of 2-Mercaptoethanol, before using an Omni Bead Ruptor Elite to disrupt the larval tissue at 4.5m/s for 30 seconds. Samples were further disrupted by passing tissue through a Qiagen QIAshredder Mini spin column before RNA extraction was performed using the Qiagen RNeasy Mini Kit according to the manufacturer’s protocol. RNA was then sent to GENEWIZ for library preparation and 150bp paired-end Illumina sequencing on the Illumina HiSeq 4000. Reads from this dataset can be found at NCBI SRA (Supplementary Table 1).

### RNA extraction, library preparation, and sequencing: dissected larval salt-secreting glands

mRNA was collected from dissected salt-secreting glands of *Ae. togoi larvae*. From hatching until RNA-extraction at the 4^th^ instar stage, four biological replicates (containing 55-65 salt-secreting glands from individual larvae each) per condition were reared in either dechlorinated Seattle tap water or dechlorinated Seattle tap water supplemented with 35 g/L sea salts (Instant-Ocean). Larvae reared in freshwater were dissected in *Ae. aeygpti* physio-logical saline solution, adjusted to pH 7.2, containing (in mmol L^-1^): 60 NaCl, 3 KCl, 5 NaHCO_3_, 0.6 MgSO_4_, 5 CaCl_2_, 25 HEPES, 10 Glucose, 5 Succinic acid, 5 Malic acid, 10 Citric acid tri-sodium salt, 3.37 L-arginine, 14.4 L-leucine, 8.74 L-histidine, 9.1 L-glutamine, and 5 L-proline (Clark and Bradley, 1996). Larvae reared in seawater (SW) were dissected in a modified *Ae. aegypti* physiological saline adjusted to pH 7.8, containing (in mmol L^-1^): 110 NaCl, 10 KCl, 5 NaHCO_3_, 0.6 MgSO_4_, 5 CaCl_2_, 25 HEPES, 10 Glucose, 5 Succinic acid, 5 Malic acid, 10 Citric acid tri-sodium salt, 3.37 L-arginine, 14.4 L-leucine, 8.74 L-histidine, 9.1 L-glutamine, and 5 L-proline adjusted to a pH of 7.8, matching levels measured for *Ae. taeniorhynchus* in seawater (Donini et al., 2006). Salt gland tissue was sonicated for 5 seconds at amplitude 10 using a sonic dismembrator (Model 50, Fisherbrand) and a 1.8-inch probe (Model CL-18, QSonica). RNA was isolated using the Direct-zol RNA MicroPrep kit (Cat #: R2060, Zymo Research) and then sent to the Northwest Genomics Center (NWGC) for quality control (Quant-iT RNA assay from Invitrogen and fragment analyzer from Advanced Analytical), library preparation (Catalogue: 20020595, TruSeq Stranded mRNA kit, Illumina), and sequencing (Illumina HiSeq 4000, 100bp paired end). Reads from this dataset can be found at NCBI SRA (Supplementary Table 1).

### Read QC and processing

HiFi reads were trimmed to remove adapter content using Lima (v2.13.0) and PCR-deduplicated with pbmarkdup (v1.2.0). HiFi, ONT, and Hi-C read quality were checked using FastQC (v0.12.1) (Andrews, 2010). Hi-C reads were trimmed to remove the front five base pairs using fastp (Chen et al., 2018). Genomescope2 (v2.0) was used to estimate heterozygosity, repeat content, and error rate of HiFi reads based on Meryl (v1.4.1) k-mer analysis (Supplementary Figure 1) (Ranallo-Benavidez et al., 2020; Rhie et al., 2020).

### Genome assembly

The primary genome assembly was performed using Hifiasm (v0.24.0) (Cheng et al., 2021). HiFi reads were used as the primary input reads while additional integration of ultra-long ONT and Hi-C reads were used to improve contiguity and haplotype resolution, respectively. The resulting assembly of primary contigs was used for further assembly steps. Purging of haplotypic duplication was done using purge_dups (v1.2.5) (Guan et al., 2020). The Arima Genomics mapping_pipeline (A160156_v03) was used to deduplicate and align Hi-C reads to the purged contig assembly. Scaffolding was performed using YaHS (v1.2.2) with default parameters (Zhou et al., 2023).

The complete mitochondrial genome sequence was assembled using MitoHifi (v3.2.2), using default parameters with The Invertebrate Mitochondrial Code (-o 5) (Uliano-Silva et al., 2023).

### Gene-order synteny between Aedes species

Syntenic blocks on analogous chromosome scaffolds between species were identified and visualized with chromsyn (v1.3.2) based on gene-order synteny of the locations of BUSCO single-copy orthologues (Edwards et al., 2022). The *Ae. albopictus* (GCA_035046485.1), *Ae. aegypti* (GCA_002204515.1), and *Ae. notoscriptus* (GCA_040801935.1) genomes were accessed from the NCBI genome database.

### Genome Assembly Curation and Finalizing

Several changes to the YaHS scaffolded assembly were made by manual curation using the Juicer, Juicebox, and 3D-DNA suite of tools. Juicer (v2.0) (Durand et al., 2016a) was used to generate the Hi-C contact map based on the scaffolded genome assembly, which was visualized and manually edited using Juicebox (v3.1.4) (Durand et al., 2016b).

Manual curation was conducted with guidance from gene-order synteny, alignment of ONT long-reads to the scaffolded *Ae. togoi* genome assembly, and Hi-C contact map signal patterns. Gene order synteny between *Ae. togoi, Ae. notoscriptus*, and *Ae. triseriatus* genomes (personal communication with John Soghigian), made it possible to identify instances of chromosome structure similarity and potential translocations and inversions (Supplementary Figure 2). *Ae. notoscriptus* and *Ae. triseriatus* were chosen to guide curation of *Ae. togoi* chromosome scaffolds, over other *Aedes* genomes, because these two species are more closely related to *Ae. togoi* and have chromosome-scale genome assemblies (Yeo et al., 2025). Syntenic blocks on *Ae. togoi* chromosome scaffolds that aligned with non-equivalent *Ae. notoscriptus* and *Ae. triseriatus* chromosome scaffolds were rearranged to agree with the *Ae. notoscriptus* and *Ae. triseriatus* chromosome structure, by breaking chromosome scaffolds and rearranging chromosome fragments where needed in Juicebox (Supplementary Figure 4). Inversions of regions within larger syntenic blocks were not edited. When a potential adjustment was identified, the sequence range for a potential scaffold cut-point was located by identifying the two BUSCO genes closest to either side of the point where the *Ae. togoi* scaffold starts to align with different scaffolds on the *Ae. notoscriptus* and *Ae. triseriatus* genomes (Supplementary Figure 2). The region between the two BUSCO genes identifies the potential sequence range for the proposed cut-site, which is determined by analyzing Hi-C contact signal patterns, looking for irregularities that suggest mis-assembly at that region. A finer-resolution cut-site is identified within the potential sequence region by aligning ONT long-reads to the scaffolded assembly and looking for gaps in read coverage caused by poor read alignment, truncated reads, or N insertions in the genome assembly (Supplementary Figure 3). In the situation of there being multiple potential cut-sites of equally low read coverage within the proposed region, the final decision was made based on the Hi-C contact signal pattern–specifically, potential cut-sites at regions with no or little signal were prioritized over potential cut-sites in regions of dense contact signal. Once an exact cut-site was identified, a 1kb region encapsulating the cut-site was removed from the scaffold and moved to debris as a separate scaffold, thereby creating a break in the existing scaffold. The resulting chromosome fragments on either side of the cut were then rearranged to match the order of syntenic blocks in the *Ae. notoscriptus* and *Ae. triseriatus* genomes. Finally, the cuts and chromosome rearrangements were accepted if the resulting Hi-C contact signal pattern no longer had signs of mis-assembly (Supplementary Figure 4). The chromosome structure of *Ae. notoscriptus* and *Ae. triseriatus* agreed for chromosomes 1 and 3, but for chromosome 2 there was a chromosomal rearrangement in *Ae. triseriatus* not found in *Ae. notoscriptus* (personal communication with John Soghigian). In this case, curation of the *Ae. togoi* chromosome 2 was guided solely by *Ae. triseriatus* chromosome structure because that suggested rearrangement was better supported by Hi-C contact signal pattern at this chromosome, while the Hi-C contact signal in the arrangement matching the *Ae. notoscriptus* chromosome 2 structure suggested mis-assembly. A total of five cuts were made: two in scaffold_1 to separate out chromosomes 1 and 3, another two in scaffold_1 to excise mitochondrial genome sequence, and one in scaffold_2 to reorder the arms of chromosome 2 (Supplementary Figure 4). 3D-DNA’s post-review pipeline was used to apply changes on the edited assembly map to the scaffolded genome assembly and for re-scaffolding, following which the assembly’s scaffolds adopt the ‘HiC_scaffold_’ naming scheme (Dudchenko et al., 2017).

Finally, stretches of N sequences (gaps) in the scaffolded assembly were closed with TGS-GapCloser (v1.2.1), using HiFi reads as input for gap-filling and Racon as the error-correction module (Xu et al., 2020).

The assembled mitochondrial genome sequence, excised from the main assembly, was identified as HiC_scaffold_219 following the re-scaffolding process. This mitochondrial genome scaffold was separated from the rest of the genome assembly.

### Genome assembly statistics

Genome assembly stats were measured using BBMap ‘stats.sh’ (v35.85) (Bushnell, 2014). Genome completeness was measured by running BUSCO (v5.5.0) (Manni et al., 2021), for conserved gene-level estimation with dipteran databases as reference.

### Repeat Content Annotation

Repeat sequence composition was estimated by using RepeatModeler (v2.0.6) to create a *de novo* predicted repeat library and RepeatMasker (v4.1.9) to search for and mask repeat sequences (Flynn et al., 2020; Tarailo-Graovac and Chen, 2009). The accuracy of this methodology was somewhat validated by estimating the repeat content of the *Ae. aegypti* genome in the same way and finding an estimated 75.30%, comparable to previous estimations of 65-78% (Matthews et al., 2018; Morinaga et al., 2025) (Supplementary Table 2; Supplementary Figure 5).

### Protein-coding gene set annotation

BRAKER3 (v3.0.8) (Gabriel et al., 2024) was used to identify protein-coding regions within the soft-masked *Ae. togoi* genome assembly, utilizing the OrthoDb v10 Arthropoda ortholog database as inputs (Kriventseva et al., 2019) and RNA-seq libraries from *Ae. togoi* adult male heads and bodies, adult female heads and bodies, whole larvae, and larval anal canals, as described above, as well as data from larval anal papillae (to be described elsewhere).

### Functional annotation

Functional information for predicted genes were drawn from GO-term and PFAM domain annotations provided by eggnog-mapper (v2.1.12) (Cantalapiedra et al., 2021; Huerta-Cepas et al., 2019). Three enrichment analyses were conducted. The first was to identify GO-terms enriched among salt-secreting gland transcripts compared to transcripts from the whole larvae transcriptome, by considering transcripts expressed in the salt-secreting gland among the top 50% with positive TPM-fold change when compared to transcripts expressed in the whole larvae in the matching freshwater or saltwater condition. The second analysis was to identify GO-terms enriched in the whole larvae when the animal is under saltwater conditions. The third analysis was to identify GO-terms enriched in the salt-secreting gland when the animal is under saltwater conditions. GO-term enrichment analyses were conducted with topGO (v2.54.0), using a weighted FDR < 0.05 cutoff for significant enrichment (Alexa and Rahnenfuhrer, 2023).

Annotation of protein coding genes, and their transcript and protein products, was done based on protein sequence similarity to *Ae. aegypti* and *Ae. camptorhynchus* proteins, two other *Aedes* genus species with previously annotated reference genomes (Matthews et al., 2018). *Ae. aegypti* (NCBI Aedes aegypti Annotation Release 101) and *Ae. camptorhynchus* (GCF_037179485.1-RS_2024_05) protein sequences were taken from NCBI Genome Annotation databases. Additional *Ae. aegypti* annotations curated through Mosquito Cell Atlas were also included (Goldman et al., 2025). First, a search for high-confidence one-to-one orthologues was conducted using Orthologr’s blast_rec function (v0.4.2) to search for reciprocal best hits (RBH) between the reference species’ proteome and the BRAKER3-predicted *Aedes togoi* protein sequences (Drost et al., 2015). Additionally, to gain a sense of protein function, albeit at a lower level of confidence, additional *Ae. togoi* proteins were annotated using a best hit (BH) BLAST approach, using the *Ae. togoi* proteins that did not find a RBH match as a query to search against the reference species’ proteome again, using an e-value threshold of 1e-6. This process was done separately for both *Ae. aegypti* and *Ae. camptorhynchus* proteomes before the results were combined. Names for *Ae. togoi* transcripts were given based on the matching *Ae. camptorhynchus* annotation name, due to higher protein sequence identity matches in general, compared to *Ae. aegypti* transcripts.

### Differential expression of whole larvae and dissected salt-secreting gland tissues

Transcript abundance from whole larval and dissected salt gland RNA datasets were separately measured using the Nextflow (v25.04.6) nf-core rnaseq pipeline (v3.22.2), which utilizes SALMON to generate count metrics for each transcript (Ewels et al., 2020; Patro et al., 2017). Transcript counts were analyzed for differential expression with DESeq2 (v1.42.1) (Love et al., 2014). Lists of differentially expressed transcripts were filtered for significance at an adjusted p-value threshold of 0.05. Volcano plots were made using EnhancedVolcano (v1.20.0), using a log2FoldChance threshold of log_2_(1.5) (Blighe et al., 2018). Transcript counts are represented as transcript per million (TPM) values in annotation tables

## References

Afifi, S., Wahedi, A. and Paluzzi, J.-P. (2023). Functional insight and cell-specific expression of the adipokinetic hormone/corazonin-related peptide in the human disease vector mosquito, Aedes aegypti. Gen. Comp. Endocrinol. 330, 114145.

Albers, M. A., & Bradley, T. J. (2011). On the evolution of saline tolerance in the larvae of mosquitoes in the genus Ochlerotatus. Physiological and Biochemical Zoology, 84(3), 258–267.

Alexa, A., Rahnenführer, J. (2025). topGO: Enrichment Analysis for Gene Ontology. doi:10.18129/B9.bioc.topGO, R package version 2.62.0, https://bioconductor.org/packages/topGO.

Andrews, S. (2010). FastQC: A Quality Control Tool for High Throughput Sequence Data.

Asakura, K. (1980). The anal portion as a salt-excreting organ in a seawater mosquito larva, Aedes togoi Theobald. J. Comp. Physiol. B 138, 59–65.

Asakura, K. (1982). Ultrastructure and chloride cytochemistry of the hind-gut epithelium in the larvae of the seawater mosquito, Aedes togoi Theobald. Arch. Histol. Jpn. 45, 167–180.

Ayala, D., Ullastres, A. and González, J. (2014). Adaptation through chromosomal inversions in Anopheles. Front. Genet. 5,.

Ayala, D., Zhang, S., Chateau, M., Fouet, C., Morlais, I., Costantini, C., Hahn, M. W. and Besansky, N. J. (2019). Association mapping desiccation resistance within chromosomal inversions in the African malaria vector Anopheles gambiae. Mol. Ecol. 28, 1333–1342.

Berdan, E. L., Flatt, T., Kozak, G. M., Lotterhos, K. E. and Wielstra, B. (2022). Genomic architecture of supergenes: connecting form and function. Philos. Trans. R. Soc. B Biol. Sci. 377, 20210192.

Berdan, E. L., Barton, N. H., Butlin, R., Charlesworth, B., Faria, R., Fragata, I., Gilbert, K. J., Jay, P., Kapun, M., Lotterhos, K. E., et al. (2023). How chromosomal inversions reorient the evolutionary process. J. Evol. Biol. 36, 1761–1782.

Beyenbach, K. W. and Piermarini, P. M. (2011). Transcellular and Para-cellular Pathways of Transepithelial Fluid Secretion in Malpighian (renal) Tubules of the Yellow Fever Mosquito Aedes aegypti. Acta Physiol. Oxf. Engl. 202, 387–407.

Beyenbach, K. W., Skaer, H., and Julian A. T. Dow (2010). The Developmental, Molecular, and Transport Biology of Malpighian Tubules. Annu. Rev. Entomol. 55, 351–374.

Bradley, T. J. (1987). Physiology of Osmoregulation in Mosquitoes. Annu. Rev. Entomol. 32, 439–462.

Cantalapiedra, C. P., Hernández-Plaza, A., Letunic, I., Bork, P. and Huerta-Cepas, J. (2021). eggNOG-mapper v2: Functional Annotation, Orthology Assignments, and Domain Prediction at the Metagenomic Scale. Mol. Biol. Evol. 38, 5825–5829.

Chasiotis, H., Ionescu, A., Misyura, L., Bui, P., Fazio, K., Wang, J., Patrick, M., Weihrauch, D. and Donini, A. (2016). An animal homolog of plant Mep/Amt transporters promotes ammonia excretion by the anal papillae of the disease vector mosquito Aedes aegypti. J. Exp. Biol. 219, 1346–1355.

Chen, Y., Veenstra, J. A., Davis, N. T. and Hagedorn, H. H. (1994). A comparative study of leucokinin-immunoreactive neurons in insects. Cell Tissue Res. 276, 69–83.

Chen, S., Zhou, Y., Chen, Y. and Gu, J. (2018). fastp: an ultra-fast all-in-one FASTQ preprocessor. Bioinformatics 34, i884–i890.

Cheng, H., Concepcion, G. T., Feng, X., Zhang, H. and Li, H. (2021). Haplotype-resolved de novo assembly using phased assembly graphs with hifiasm. Nat. Methods 18, 170–175.

Choi, J. W. and Choi, K. S. (2024). Effect of salinity on the oviposition and growth of Ochlerotatus togoi. Ecol. Evol. 14, e11289.

Clark, T. M. and Bradley, T. J. (1996). Stimulation of Malpighian tubules from larval Aedes aegypti by secretagogues. J. Insect Physiol. 42, 593–602.

Daron, J., Bergman, A. and Lambrechts, L. (2025). Dynamics and evolution of transposable elements in mosquito genomes. Curr. Opin. Insect Sci. 71, 101406.

Derilus, D., Weedall, G. D., Vandewege, M. W., Batra, D., Sheth, M., Rowe, L. A., Escalante, A. A., Lenhart, A. and Impoinvil, L. M. (2025). Chromosome-scale genome assembly and annotation of two geographically distinct strains of malaria vector Anopheles albimanus. Sci. Rep. 15, 19448.

Donini, A., Patrick, M. L., Bijelic, G., Christensen, R. J., Ianowski, J. P., Rheault, M. R. and O’Donnell, M. J. (2006). Secretion of Water and Ions by Malpighian Tubules of Larval Mosquitoes: Effects of Diuretic Factors, Second Messengers, and Salinity. Physiol. Biochem. Zool. 79, 645–655.

Drost, H.-G., Gabel, A., Grosse, I. and Quint, M. (2015). Evidence for active maintenance of phylotranscriptomic hourglass patterns in animal and plant embryogenesis. Mol. Biol. Evol. 32, 1221–1231.

Duca, O. D., Nasirian, A., Galperin, V. and Donini, A. (2011). Pharmacological characterisation of apical Na+ and Cl– transport mechanisms of the anal papillae in the larval mosquito Aedes aegypti. J. Exp. Biol. 214, 3992–3999.

Dudchenko, O., Batra, S. S., Omer, A. D., Nyquist, S. K., Hoeger, M., Durand, N. C., Shamim, M. S., Machol, I., Lander, E. S., Aiden, A. P., et al. (2017). De novo assembly of the Aedes aegypti genome using Hi-C yields chromosome-length scaffolds. Science 356, 92–95.

Durand, N. C., Shamim, M. S., Machol, I., Rao, S. S. P., Huntley, M. H., Lander, E. S. and Aiden, E. L. (2016a). Juicer Provides a One-Click System for Analyzing Loop-Resolution Hi-C Experiments. Cell Syst. 3, 95–98.

Durand, N. C., Robinson, J. T., Shamim, M. S., Machol, I., Mesirov, J. P., Lander, E. S. and Aiden, E. L. (2016b). Juicebox Provides a Visualization System for Hi-C Contact Maps with Unlimited Zoom. Cell Syst. 3, 99–101.

Durant, A. C., Grieco Guardian, E., Kolosov, D. and Donini, A. (2021). The transcriptome of anal papillae of Aedes aegypti reveals their importance in xenobiotic detoxification and adds significant knowledge on ion, water and ammonia transport mechanisms. J. Insect Physiol. 132, 104269.

Edwards, R. J., Dong, C., Park, R. F. and Tobias, P. A. (2022). A phased chromosome-level genome and full mitochondrial sequence for the dikaryotic myrtle rust pathogen, Austropuccinia psidii. 2022.04.22.489119.

Ewels, P. A., Peltzer, A., Fillinger, S., Patel, H., Alneberg, J., Wilm, A., Garcia, M. U., Di Tommaso, P. and Nahnsen, S. (2020). The nf-core framework for community-curated bioinformatics pipelines. Nat. Biotechnol. 38, 276–278.

Feder, J. L. and Nosil, P. (2009). CHROMOSOMAL INVERSIONS AND SPECIES DIFFERENCES: WHEN ARE GENES AFFECTING ADAPTIVE DIVERGENCE AND REPRODUCTIVE ISOLATION EXPECTED TO RESIDE WITHIN INVERSIONS? Evolution 63, 3061–3075.

Flynn, J. M., Hubley, R., Goubert, C., Rosen, J., Clark, A. G., Feschotte, C. and Smit, A. F. (2020). RepeatModeler2 for automated genomic discovery of transposable element families. Proc. Natl. Acad. Sci. 117, 9451–9457.

Fontaine, M. C., Pease, J. B., Steele, A., Waterhouse, R. M., Neafsey, D. E., Sharakhov, I. V., Jiang, X., Hall, A. B., Catteruccia, F., Kakani, E., et al. (2015). Extensive introgression in a malaria vector species complex revealed by phylogenomics. Science 347, 1258524.

Gabriel, L., Brůna, T., Hoff, K. J., Ebel, M., Lomsadze, A., Borodovsky, M. and Stanke, M. (2024). BRAKER3: Fully automated genome annotation using RNA-seq and protein evidence with GeneMark-ETP, AUGUSTUS, and TSEBRA. Genome Res. 34, 769–777.

Gade, G., Hoffmann, K. H. and Spring, J. H. (1997). Hormonal regulation in insects: facts, gaps, and future directions. Physiol. Rev. 77, 963–1032.

Ghosh, S. K., Podder, D., Panja, A. K. and Mukherjee, S. (2020). In target areas where human mosquito-borne diseases are diagnosed, the inclusion of the pre-adult mosquito aquatic niches parameters will improve the integrated mosquito control program. PLoS Negl. Trop. Dis. 14, e0008605.

Goldman, O. V., DeFoe, A. E., Qi, Y., Jiao, Y., Weng, S.-C., Houri-Zeevi, L., Lakhiani, P., Morita, T., Razzauti, J., Rosas-Villegas, A., et al. (2025). Mosquito Cell Atlas: A single-nucleus transcriptomic atlas of the adult Aedes aegypti mosquito.

Guan, D., McCarthy, S. A., Wood, J., Howe, K., Wang, Y. and Durbin, R. (2020). Identifying and removing haplotypic duplication in primary genome assemblies. Bioinformatics 36, 2896–2898.

Habtewold, T., Wagah, M., Tambwe, M. M., Moore, S., Windbichler, N., Christophides, G., Johnson, H., Heaton, H., Collins, J., Krasheninnikova, K., et al. (2024). A chromosomal reference genome sequence for the malaria mosquito, Anopheles gambiae, Giles, 1902, Ifakara strain. Wellcome Open Res. 8, 74.

Hansen, K. K., Stafflinger, E., Schneider, M., Hauser, F., Cazzamali, G., Williamson, M., Kollmann, M., Schachtner, J. and Grimme-likhuijzen, C. J. P. (2010). Discovery of a Novel Insect Neuropeptide Signaling System Closely Related to the Insect Adipokinetic Hormone and Corazonin Hormonal Systems. J. Biol. Chem. 285, 10736–10747.

Hesson, J. C., Haba, Y., McBride, C. S., Sheerin, E., Mathers, T. C., Paulini, M., Pointon, D.-L. B., Torrance, J. W., Sadasivan Baby, C., Wood, J. M. D., et al. (2025). A chromosomal reference genome sequence for the northern house mosquito, Culex pipiens form pipiens, Linnaeus, 1758. Wellcome Open Res. 10, 107.

Hoffmann, A. A., Sgrò, C. M. and Weeks, A. R. (2004). Chromosomal inversion polymorphisms and adaptation. Trends Ecol. Evol. 19, 482–488.

Horváth, V., Merenciano, M. and González, J. (2017). Revisiting the Relationship between Transposable Elements and the Eukaryotic Stress Response. Trends Genet. 33, 832–841.

Huerta-Cepas, J., Szklarczyk, D., Heller, D., Hernández-Plaza, A., Forslund, S. K., Cook, H., Mende, D. R., Letunic, I., Rattei, T., Jensen, L. J., et al. (2019). eggNOG 5.0: a hierarchical, functionally and phylogenetically annotated orthology resource based on 5090 organisms and 2502 viruses. Nucleic Acids Res. 47, D309–D314.

Ingham, V. A., Tennessen, J. A., Lucas, E. R., Elg, S., Yates, H. C., Carson, J., Guelbeogo, W. M., Sagnon, N., Hughes, G. L., Heinz, E., et al. (2021). Integration of whole genome sequencing and transcriptomics reveals a complex picture of the reestablishment of insecticide resistance in the major malaria vector Anopheles coluzzii. PLOS Genet. 17, e1009970.

Iv, W. C. B. and Rai, K. S. (1988). Genome evolution in mosquitoes: intraspecific and interspecific variation in repetitive DNA amounts and organization. Genet. Res. 51, 185–196.

Jay, P. and Joron, M. (2022). The double game of chromosomal inversions in a neotropical butterfly. C. R. Biol. 345, 57–73.

Jay, P., Whibley, A., Frézal, L., Cara, M. Á. R. de, Nowell, R. W., Mallet, J., Dasmahapatra, K. K. and Joron, M. (2018). Supergene Evolution Triggered by the Introgression of a Chromosomal Inversion. Curr. Biol. 28, 1839–1845.e3.

Jonusaite, S. and Himmerkus, N. (2024). Paracellular barriers: Advances in assessing their contribution to renal epithelial function. Comp. Biochem. Physiol. A. Mol. Integr. Physiol. 298, 111741.

Jonusaite, S., Kelly, S. P., & Donini, A. (2016). The response of claudin-like transmembrane septate junction proteins to altered environmental ion levels in the larval mosquito Aedes aegypti. Journal of Comparative Physiology B, 186(5), 589–602.

Kandel, Y., Pinch, M., Lamsal, M., Martinez, N. and Hansen, I. A. (2022). Exploratory phosphoproteomics profiling of Aedes aegypti Malpighian tubules during blood meal processing reveals dramatic transition in function. PLOS ONE 17, e0271248.

Kengne, P., Charmantier, G., Blondeau-Bidet, E., Costantini, C. and Ayala, D. (2019). Tolerance of disease-vector mosquitoes to brackish water and their osmoregulatory ability. Ecosphere 10, e02783.

Kent, T. V., Uzunović, J. and Wright, S. I. (2017). Coevolution between transposable elements and recombination. Philos. Trans. R. Soc. B Biol. Sci. 372, 20160458.

Kriventseva, E. V., Kuznetsov, D., Tegenfeldt, F., Manni, M., Dias, R., Simão, F. A. and Zdobnov, E. M. (2019). OrthoDB v10: sampling the diversity of animal, plant, fungal, protist, bacterial and viral genomes for evolutionary and functional annotations of orthologs. Nucleic Acids Res. 47, D807–D811.

Kubrak, O. I., Lushchak, O. V., Zandawala, M. and Nässel, D. R. (2016). Systemic corazonin signalling modulates stress responses and metabolism in Drosophila. Open Biol. 6, 160152.

Liang, J., Rose, N. H., Brusentsov, I. I., Lukyanchikova, V., Karagodin, D. A., Feng, Y., Yurchenko, A. A., Sharma, A., Sylla, M., Lutomiah, J., et al. (2025). Chromosomal Inversions and Their Potential Impact on the Evolution of Arboviral Vector Aedes aegypti. Genome Biol. Evol. 17, evaf118.

Liu, W., Cheng, P., An, S., Zhang, K., Gong, M., Zhang, Z. and Zhang, R. (2023). Chromosome-level assembly of Culex pipiens molestus and improved reference genome of Culex pipiens pallens (Culicidae, Diptera). Mol. Ecol. Resour. 23, 486–498.

Love, M. I., Huber, W. and Anders, S. (2014). Moderated estimation of fold change and dispersion for RNA-seq data with DESeq2. Genome Biol. 15, 550.

Love, R. R., Redmond, S. N., Pombi, M., Caputo, B., Petrarca, V., della Torre, A., The Anopheles gambiae 1000 Genomes Consortium and Besansky, N. J. (2019). In Silico Karyotyping of Chromosomally Polymorphic Malaria Mosquitoes in the Anopheles gambiae Complex. G3 GenesGenomesGenetics 9, 3249–3262.

Lushasi, S. C., Mwalugelo, Y. A., Swai, J. K., Mmbando, A. S., Muyaga, L. L., Nyolobi, N. K., Mutashobya, A., Mmbaga A. T., Kunambi, H. J., Twaha, S., Mwema, M. F., and Lwetoijera, D. W. (2024). The interspecific competition between larvae of Aedes aegypti and major African malaria vectors in a semi-field system in Tanzania. Insects, 16(1), 34.

Ma, L.-J., Cao, L.-J., Chen, J.-C., Tang, M.-Q., Song, W., Yang, F.-Y., Shen, X.-J., Ren, Y.-J., Yang, Q., Li, H., et al. (2024). Rapid and Repeated Climate Adaptation Involving Chromosome Inversions following Invasion of an Insect. Mol. Biol. Evol. 41, msae044.

Main, B. J., Lee, Y., Ferguson, H. M., Kreppel, K. S., Kihonda, A., Govella, N. J., Collier, T. C., Cornel, A. J., Eskin, E., Kang, E. Y., et al. (2016). The Genetic Basis of Host Preference and Resting Behavior in the Major African Malaria Vector, Anopheles arabiensis. PLOS Genet. 12, e1006303.

Manni, M., Berkeley, M. R., Seppey, M., Simão, F. A. and Zdobnov, E. M. (2021). BUSCO Update: Novel and Streamlined Workflows along with Broader and Deeper Phylogenetic Coverage for Scoring of Eukaryotic, Prokaryotic, and Viral Genomes. Mol. Biol. Evol. 38, 4647–4654.

Matthews, B. J., Dudchenko, O., Kingan, S. B., Koren, S., Antoshechkin, I., Crawford, J. E., Glassford, W. J., Herre, M., Redmond, S. N., Rose, N. H., et al. (2018). Improved reference genome of Aedes aegypti informs arbovirus vector control. Nature 563, 501–507.

McGinnis, K. M. and Brust, R. A. (1983). Effect of Different Sea Salt Concentrations and Temperatures on Larval Development of Aedes togoi (Diptera: Culicidae) from British Columbia. Environ. Entomol. 12, 1406–1411.

McLain, D. K., Rai, K. S. and Fraser, M. J. (1987). Intraspecific and interspecific variation in the sequence and abundance of highly repeated DNA among mosquitoes of the Aedes albopictus subgroup. Heredity 58, 373–381.

Melo, E. S. de and Wallau, G. L. (2020). Mosquito genomes are frequently invaded by transposable elements through horizontal transfer. PLOS Genet. 16, e1008946.

Morinaga, G., Balcazar, D., Badolo, A., Iyaloo, D., Tantely, M. L., Mouillaud, T., Sharakhova, M., Geib, S. M., Paupy, C., Ayala, D., et al. (2025). From Macro to Micro: De Novo Genomes of Aedes Mosquitoes Enable Comparative Genomics Among Close and Distant Relatives. Genome Biol. Evol. 17, evaf142.

Mugenzi, L. M. J., Tekoh, T. A., Ntadoun, S. T., Chi, A. D., Gadji, M., Menze, B. D., Tchouakui, M., Irving, H., Wondji, M. J., Weedall, G. D., et al. (2024). Association of a rapidly selected 4.3kb transposon-containing structural variation with a P450-based resistance to pyrethroids in the African malaria vector Anopheles funestus. PLOS Genet. 20, e1011344.

Noden, B. H., O’NEAL, P. A., Fader, J. E., & Juliano, S. A. (2016). Impact of inter-and intra-specific competition among larvae on larval, adult, and life-table traits of A edes aegypti and A edes albopictus females. Ecological entomology, 41(2), 192–200.

Palatini, U., Masri, R. A., Cosme, L. V., Koren, S., Thibaud-Nissen, F., Biedler, J. K., Krsticevic, F., Johnston, J. S., Halbach, R., Crawford, J. E., et al. (2020). Improved reference genome of the arboviral vector Aedes albopictus. Genome Biol. 21, 215.

Patrick, M. L., & Bradley, T. J. (2000). The physiology of salinity tolerance in larvae of two species of Culex mosquitoes: the role of compatible solutes. Journal of Experimental Biology, 203(4), 821–830.

Patrick, M. L., Donini, A., Zogby, A., Morales, C., O’Donnell, M. J. and Gill, S. S. (2024). Proton-driven sodium secretion in a saline water animal. Sci. Rep. 14, 12738.

Patro, R., Duggal, G., Love, M. I., Irizarry, R. A. and Kingsford, C. (2017). Salmon provides fast and bias-aware quantification of transcript expression. Nat. Methods 14, 417–419.

Peach, D. A. H. and Matthews, B. J. (2020). Modeling the Putative Ancient Distribution of Aedes togoi (Diptera: Culicidae). J. Insect Sci. 20, 7.

Phelan, O. (2023). An investigation into the salinity tolerance, oviposition behavior and breeding habitat ecology of a widespread (Aedes aegypti) and restricted (Aedes togoi) mosquito species (T). University of British Columbia. Retrieved from https://open.library.ubc.ca/collections/ub-ctheses/24/items/1.0434287

Piermarini, P. M., Hine, R. M., Schepel, M., Miyauchi, J. and Beyenbach, K. W. (2011). Role of an apical K,Cl cotransporter in urine formation by renal tubules of the yellow fever mosquito (Aedes aegypti). Am. J. Physiol. Regul. Integr. Comp. Physiol. 301, R1318–1337.

Piermarini, P. M., Dunemann, S. M., Rouhier, M. F., Calkins, T. L., Raphemot, R., Denton, J. S., Hine, R. M. and Beyenbach, K. W. (2015). Localization and role of inward rectifier K+ channels in Malpighian tubules of the yellow fever mosquito Aedes aegypti. Insect Biochem. Mol. Biol. 67, 59–73.

Piermarini, P. M., Akuma, D. C., Crow, J. C., Jamil, T. L., Kerkhoff, W. G., Viel, K. C. M. F. and Gillen, C. M. (2017). Differential expression of putative sodium-dependent cation-chloride cotransporters in Aedes aegypti. Comp. Biochem. Physiol. A. Mol. Integr. Physiol. 214, 40–49.

Ranallo-Benavidez, T. R., Jaron, K. S. and Schatz, M. C. (2020). GenomeScope 2.0 and Smudgeplot for reference-free profiling of polyploid genomes. Nat. Commun. 11, 1432.

Rheault, M. R., Okech, B. A., Keen, S. B., Miller, M. M., Meleshkevitch, E. A., Linser, P. J., Boudko, D. Y., and Harvey, W. R. (2007). Molecular cloning, phylogeny and localization of AgNHA1: the first Na+/H+ antiporter (NHA) from a metazoan, Anopheles gambiae. Journal of Experimental Biology, 210(21), 3848–3861.

Rhie, A., Walenz, B. P., Koren, S. and Phillippy, A. M. (2020). Merqury: reference-free quality, completeness, and phasing assessment for genome assemblies. Genome Biol. 21, 245.

Rosen, L., Tesh, R. B., Lien, J. C. and Cross, J. H. (1978). Transovarial Transmission of Japanese Encephalitis Virus by Mosquitoes. Science 199, 909–911.

Ryazansky, S. S., Chen, C., Potters, M., Naumenko, A. N., Lukyanchikova, V., Masri, R. A., Brusentsov, I. I., Karagodin, D. A., Yurchenko, A. A., dos Anjos, V. L., et al. (2024a). The chromo-some-scale genome assembly for the West Nile vector Culex quinquefasciatus uncovers patterns of genome evolution in mosquitoes. BMC Biol. 22, 16.

Ryazansky, S. S., Chen, C., Potters, M., Naumenko, A. N., Lukyanchikova, V., Masri, R. A., Brusentsov, I. I., Karagodin, D. A., Yurchenko, A. A., dos Anjos, V. L., et al. (2024b). The chromo-some-scale genome assembly for the West Nile vector Culex quinquefasciatus uncovers patterns of genome evolution in mosquitoes. BMC Biol. 22, 16.

Sajadi, F., Uyuklu, A., Paputsis, C., Lajevardi, A., Wahedi, A., Ber, L. T., Matei, A. and Paluzzi, J.-P. V. (2020). CAPA neuropeptides and their receptor form an anti-diuretic hormone signaling system in the human disease vector, Aedes aegypti. Sci. Rep. 10, 1755.

Sajadi, F., Vergara-Martínez, M. F., & Paluzzi, J. P. V. (2023). The V-type H+-ATPase is targeted in antidiuretic hormone control of the Malpighian “renal” tubules. Proceedings of the National Academy of Sciences, 120(51), e2308602120.

Schrader, L. and Schmitz, J. (2019). The impact of transposable elements in adaptive evolution. Mol. Ecol. 28, 1537–1549.

Sharakhova, M. V. and Sharakhov, I. V. (2025). Chromosomal rearrangements in mosquitoes: from micro-to macroevolution. Curr. Opin. Insect Sci. 71, 101393.

Small, S. T., Costantini, C., Sagnon, N., Guelbeogo, M. W., Emrich, S. J., Kern, A. D., Fontaine, M. C. and Besansky, N. J. (2023). Standing genetic variation and chromosome differences drove rapid ecotype formation in a major malaria mosquito. Proc. Natl. Acad. Sci. 120, e2219835120.

Smith, K. E., VanEkeris, L. A., Okech, B. A., Harvey, W. R. and Linser, P. J. (2008). Larval anopheline mosquito recta exhibit a dramatic change in localization patterns of ion transport proteins in response to shifting salinity: a comparison between anopheline and culicine larvae. J. Exp. Biol. 211, 3067–3076.

Stapley, J., Santure, A. W. and Dennis, S. R. (2015). Transposable elements as agents of rapid adaptation may explain the genetic paradox of invasive species. Mol. Ecol. 24, 2241–2252.

Stump, A. D., Pombi, M., Goeddel, L., Ribeiro, J. M. C., Wilder, J. A., Torre, A. D. and Besansky, N. J. (2007). Genetic exchange in 2La inversion heterokaryotypes of Anopheles gambiae. Insect Mol. Biol. 16, 703–709.

Tahami, M. S., Vargas-Chavez, C., Poikela, N., Coronado-Zamora, M., González, J. and Kankare, M. (2024). Transposable elements in a cold-tolerant fly species, Drosophila montana: a link to adaptation to the harsh cold environments. 2024.04.17.589934.

Tarailo-Graovac, M. and Chen, N. (2009). Using RepeatMasker to identify repetitive elements in genomic sequences. Curr. Protoc. Bioinforma. Chapter 4, 4.10.1-4.10.14.

Thomson, R. E. (1981) Oceanography of the British Columbia Coast. Sidney, BC. Available at: https://publications.gc.ca/collections/collection_2016/mpo-dfo/Fs41-31-56-eng.pdf.

Trimble, R. M. and Wellington, W. G. (1979). Effects of salinity on site selection by ovipositing Aedes togoi (Diptera: Culicidae). Can. J. Zool. 57, 593–596.

Uliano-Silva, M., Ferreira, J. G. R. N., Krasheninnikova, K., Darwin Tree of Life Consortium, Blaxter, M., Mieszkowska, N., Hall, N., Holland, P., Durbin, R., Richards, T., et al. (2023). MitoHiFi: a python pipeline for mitochondrial genome assembly from PacBio high fidelity reads. BMC Bioinformatics 24, 288.

Wahedi, A. and Paluzzi, J.-P. (2018). Molecular identification, transcript expression, and functional deorphanization of the adipokinetic hormone/corazonin-related peptide receptor in the disease vector, Aedes aegypti. Sci. Rep. 8, 2146.

Weetman, D., Wilding, C. S., Neafsey, D. E., Müller, P., Ochomo, E., Isaacs, A. T., Steen, K., Rippon, E. J., Morgan, J. C., Mawejje, H. D., et al. (2018). Candidate-gene based GWAS identifies reproducible DNA markers for metabolic pyrethroid resistance from standing genetic variation in East African Anopheles gambiae. Sci. Rep. 8, 2920.

Wigglesworth, V. B. (1933). The Function of the Anal Gills of the Mosquito Larva. J. Exp. Biol. 10, 16–26.

Wingett, S., Ewels, P., Furlan-Magaril, M., Nagano, T., Schoenfelder, S., Fraser, P. and Andrews, S. (2015). HiCUP: pipeline for mapping and processing Hi-C data. F1000Research 4, 1310.

Xu, M., Guo, L., Gu, S., Wang, O., Zhang, R., Peters, B. A., Fan, G., Liu, X., Xu, X., Deng, L., et al. (2020). TGS-GapCloser: A fast and accurate gap closer for large genomes with low coverage of error-prone long reads. GigaScience 9,.

Yeo, H., Chan, C., Morinaga, G., Wiegmann, B. M. and Soghigian, J. (2025). A Way Through the Trees: Molecular Phylogenies Consistently Recover Two Clades of Aedes Mosquitoes. 2025.07.16.665186.

Zamyatin, A., Avdeyev, P., Liang, J., Sharma, A., Chen, C., Lukyanchikova, V., Alexeev, N., Tu, Z., Alekseyev, M. A. and Sharakhov, I. V. (2021). Chromosome-level genome assemblies of the malaria vectors Anopheles coluzzii and Anopheles arabiensis. Giga-Science 10, giab017.

Zandawala, M., Nguyen, T., Segura, M. B., Johard, H. A. D., Amcoff, M., Wegener, C., Paluzzi, J.-P. and Nässel, D. R. (2021). A neuroendocrine pathway modulating osmotic stress in Drosophila. PLOS Genet. 17, e1009425.

Zhou, C., McCarthy, S. A. and Durbin, R. (2023). YaHS: yet another Hi-C scaffolding tool. Bioinformatics 39, btac808.

