## Supplemental Figure 1 for "Living by the sea: chromosome-scale genome assembly and salt gland transcriptomes provide insights into ion regulatory mechanisms in the saline-tolerant mosquito *Aedes togoi*"

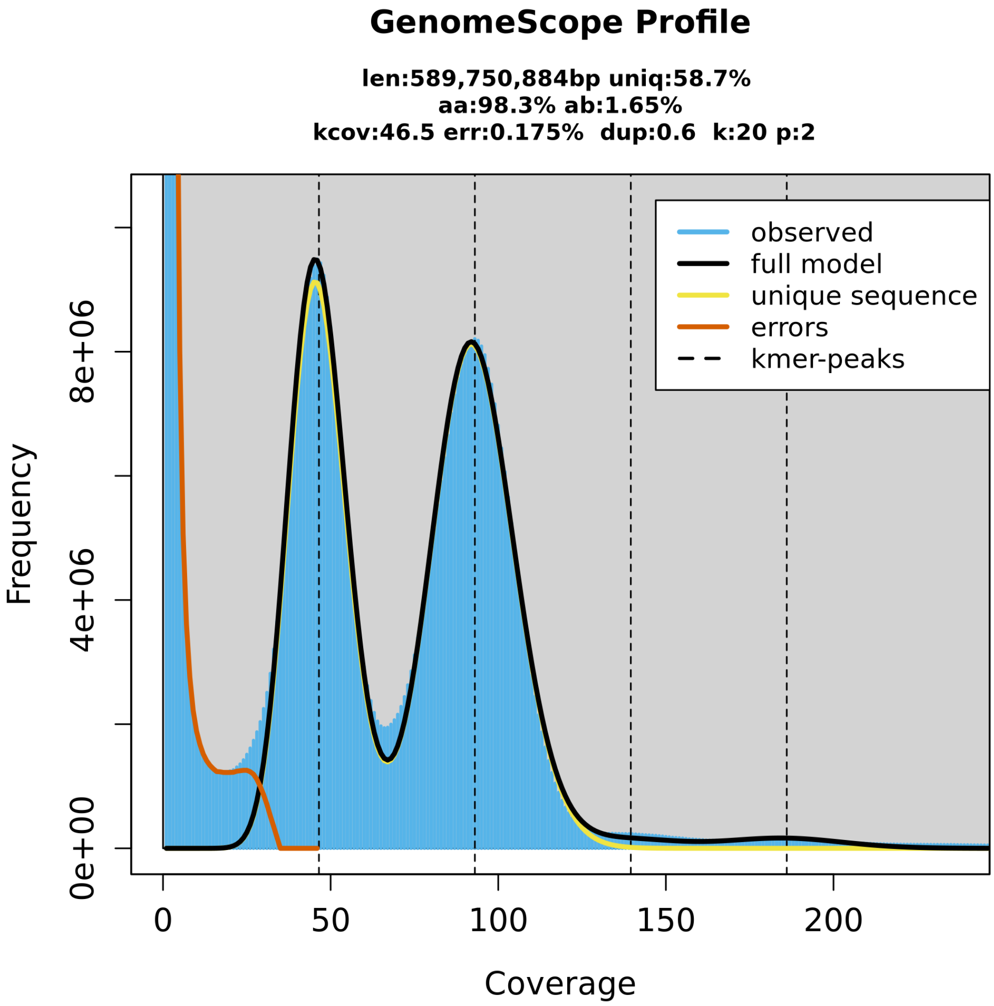


**Supplementary Figure 1. HiFi read k-mer-based estimate of genome characteristics**
*Ae. togoi* genome characteristics were estimated based on our HiFi read k-mer counts. Genomescope2 was used to estimate genome length, heterozygosity, repeat content, and HiFi read sequencing error rate were estimated based on our deduplicated HiFi reads.
