## Supplemental Figure 2 for "Living by the sea: chromosome-scale genome assembly and salt gland transcriptomes provide insights into ion regulatory mechanisms in the saline-tolerant mosquito *Aedes togoi*"

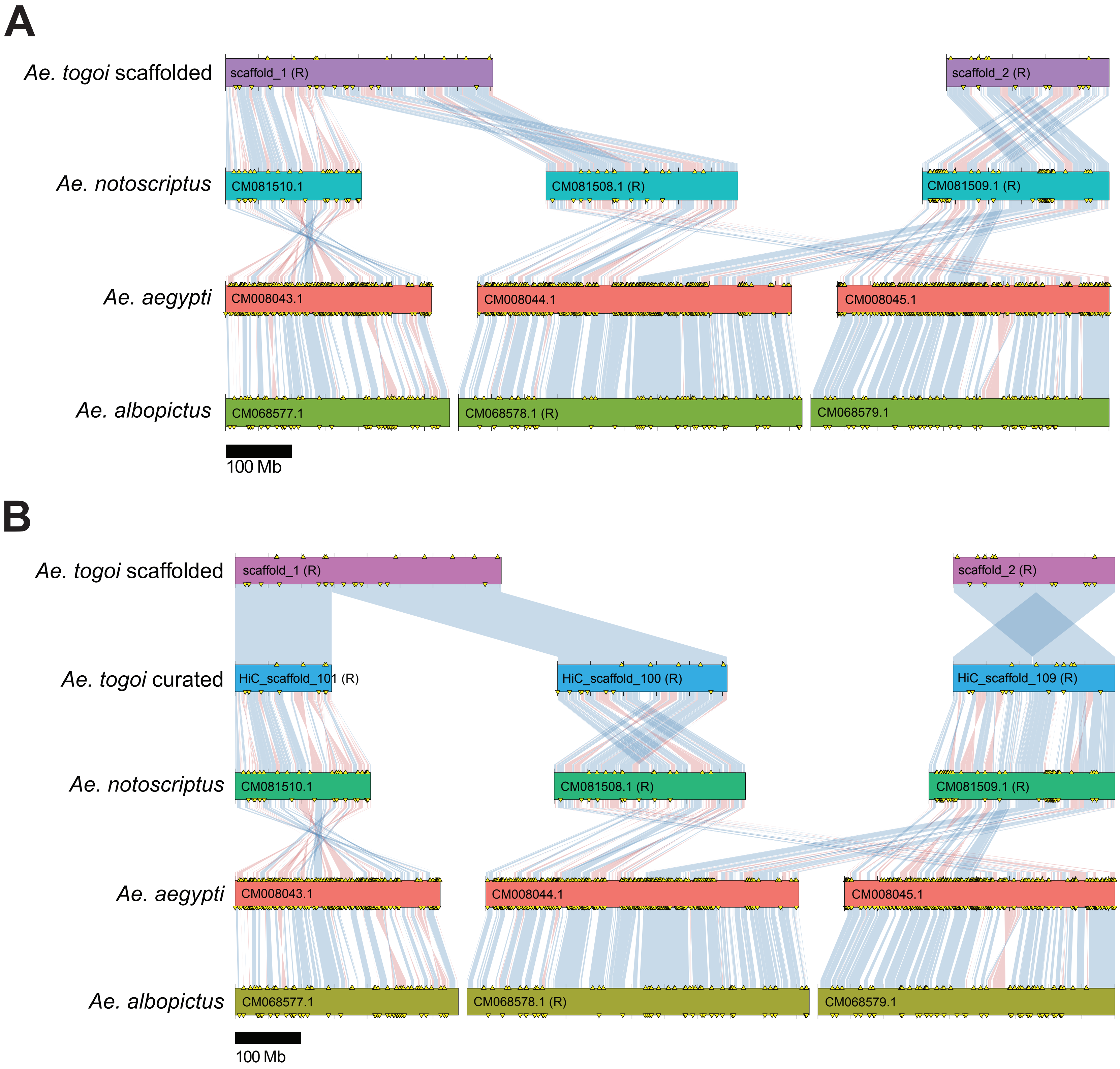


**Supplementary Figure 2. Chromosome structure of the scaffolded and curated *Aedes togoi* genome assemblies.**
(A) Chromosomal synteny between the scaffolded (no manual curation) *Ae. togoi*, *Ae. notoscriptus*, *Ae. aegypti*, and *Ae. albopictus* genomes based on the location of orthologous BUSCO genes. (B) Chromosomal synteny between the curated *Ae. togoi*, *Ae. notoscriptus*, *Ae. aegypti*, and *Ae. albopictus* genomes, highlighting changes made to chromosome structure in curation process. Chromosomal inversions are highlighted in red while syntenic blocks with matching orientation are denoted in blue. Chromosomes with the majority of synteny on the opposite strand are labelled with ‘(R)’. Plots made with chromsyn.
