## Supplemental Figure 3 for "Living by the sea: chromosome-scale genome assembly and salt gland transcriptomes provide insights into ion regulatory mechanisms in the saline-tolerant mosquito *Aedes togoi*"

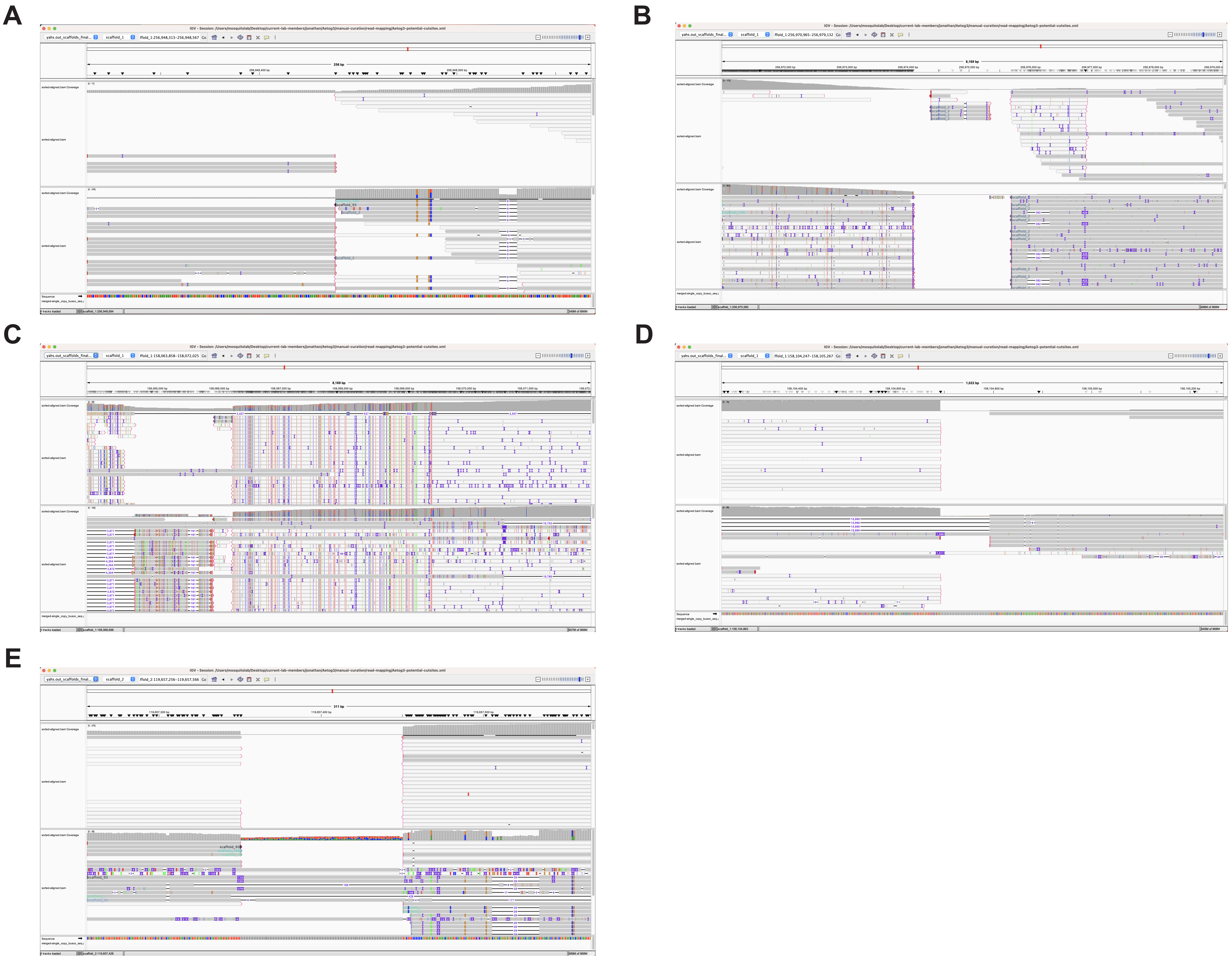


**Supplementary Figure 3. Cutsite selection in manual curation process of *Aedes togoi* genome assembly.**
IGV was used to visualize long-read sequences aligned to the scaffolded *Ae. togoi* assembly, get a fine-resolution analysis of potential regions of misassembly, and identify suitable 'cut-sites' where the scaffold would be cut into fragments to be rearranged. Cut-sites were identified at regions where there was low read coverage or gaps in read alignment. (A,B) Regions around cuts (1.1 and 1.2) in scaffold_1 made to separate chromosomes 1 and 2. The region in-between these two cut-sites was discarded to debris as a separate contig due to having high repetitive content and no reads overlapping to either side of cut-sites. (C,D) Regions around cuts (3.1 and 3.2) in scaffold_1 made to excise mitochondrial genome sequence. The region in-between these two cut-sites was separated and removed as an intact assembly of the mitochondrial genome. (E) Region around cut (2.1) in scaffold_2 made to re-order the arms of chromosome 3.
