## Supplemental Figure 4 for "Living by the sea: chromosome-scale genome assembly and salt gland transcriptomes provide insights into ion regulatory mechanisms in the saline-tolerant mosquito *Aedes togoi*"

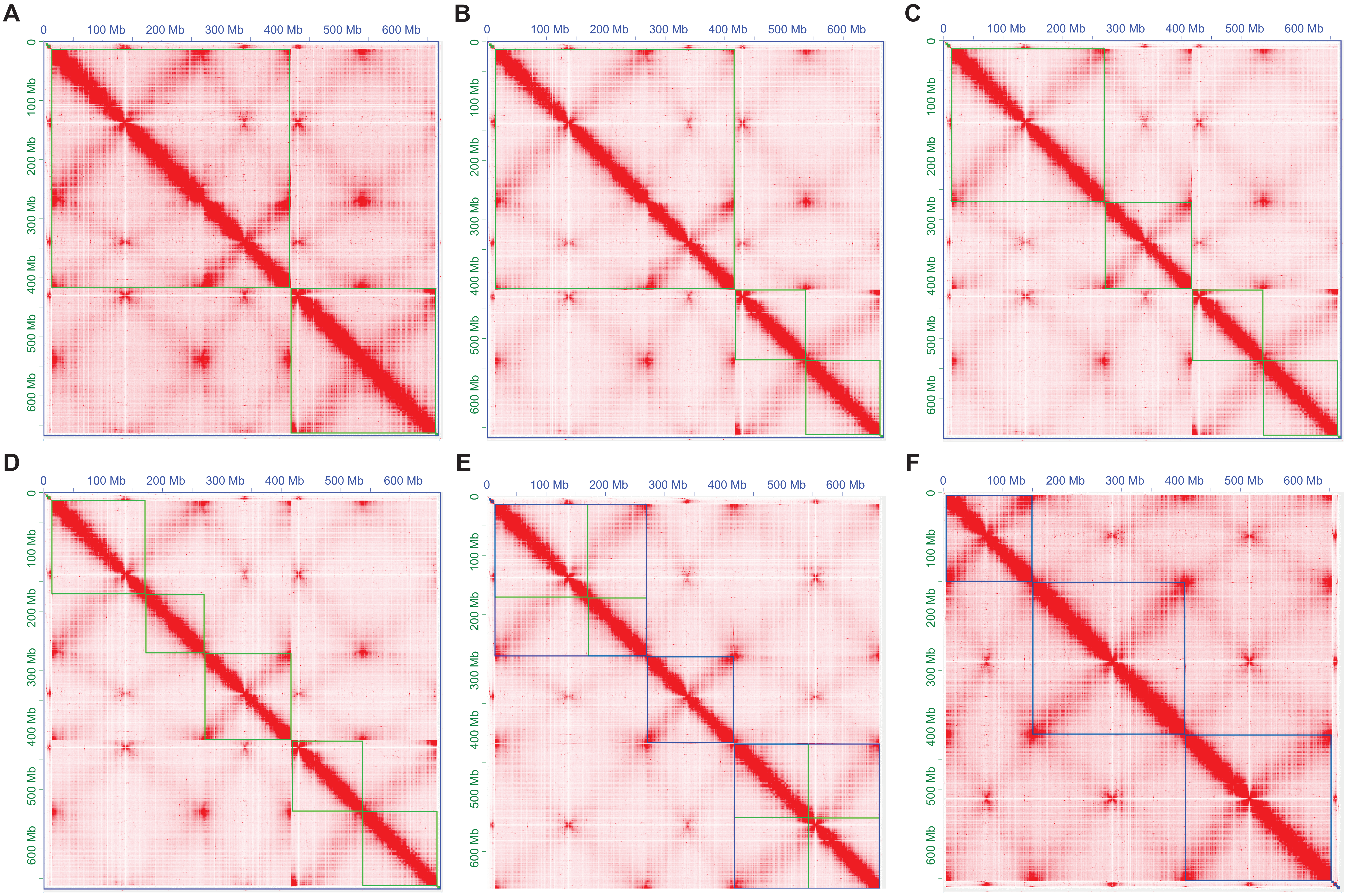


**Supplementary Figure 4. Manual curation process of *Aedes togoi* genome assembly in JuiceBox.**
Edits to the scaffolded *Ae. togoi* assembly were implemented manually in JuiceBox, to fix points of mis-assembly and achieve correct chromosome structure guided by gene-order synteny to related *Aedes* species. (A-F) Hi-C contact maps showing Hi-C reads alignment signal (in red) mapped to the genome assembly, plotted against itself on both axes. Green boundaries represent scaffold/contigs while blue boundaries represent chromosome scaffolds. (A) Contact map showing the scaffolded *Ae. togoi* genome assembly, without any edits yet. (B) Contact map showing cut-site in the centromeric region of scaffold_2. (C) Contact map showing cut-site in scaffold_1, made to separate chromosomes 1 and 2. (D) Contact map showing cut-site in scaffold_1, made to separate mitochondrial genome sequence from the main assembly of nuclear genomic sequence. (E) Contact map showing signal patterning of assembly following swapping the arms of chromosome 3 (scaffold_2), revealing a more signal pattern more consistent with correctly assembled mosquito chromosomes. (F) Contact map showing the patterning of the curated *Ae. togoi* assembly, following processing in 3D-DNA's post-review pipeline.
