## Supplemental Figure 5 for "Living by the sea: chromosome-scale genome assembly and salt gland transcriptomes provide insights into ion regulatory mechanisms in the saline-tolerant mosquito *Aedes togoi*"

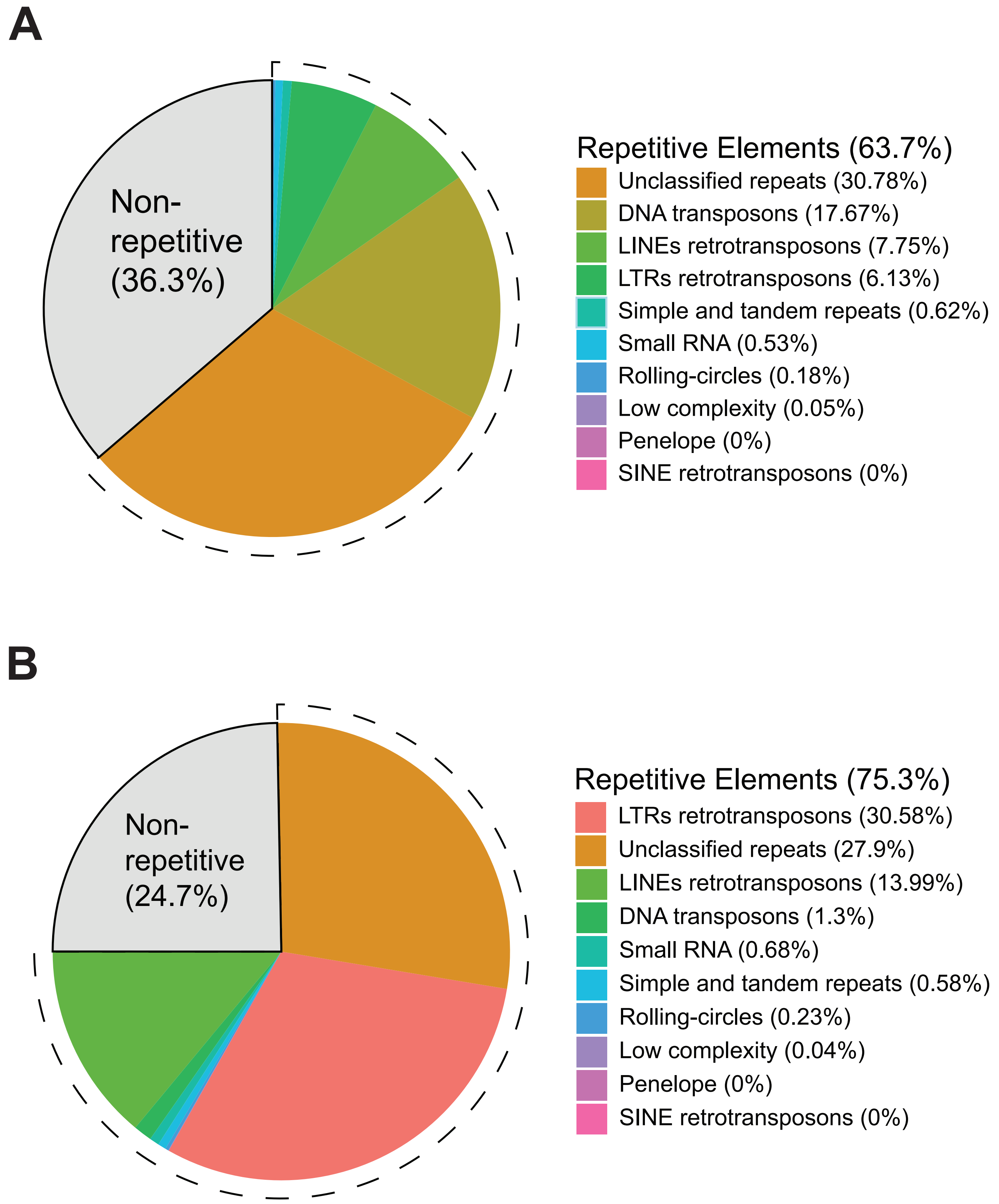


**Supplementary Figure 5. Repetitive element composition estimates by RepeatModeler and RepeatMasker.**
(A) Repetitive element composition of *Aedes togoi* genome, using *de novo* repetitive element library predicted by RepeatModeler. Repetitive elements make up 63.7% of *Ae. togoi* genomic sequence.
(B) Repetitive element composition of *Aedes aegypti* genome, using the same methodology as a way to validate *Ae. togoi* repetitive element composition results. *Ae. aegypti* genome is estimated to be composed of 75.3% repetitive sequence, comparable to estimations in previous studies.
