## Supplemental Figure 6 for "Living by the sea: chromosome-scale genome assembly and salt gland transcriptomes provide insights into ion regulatory mechanisms in the saline-tolerant mosquito *Aedes togoi*"

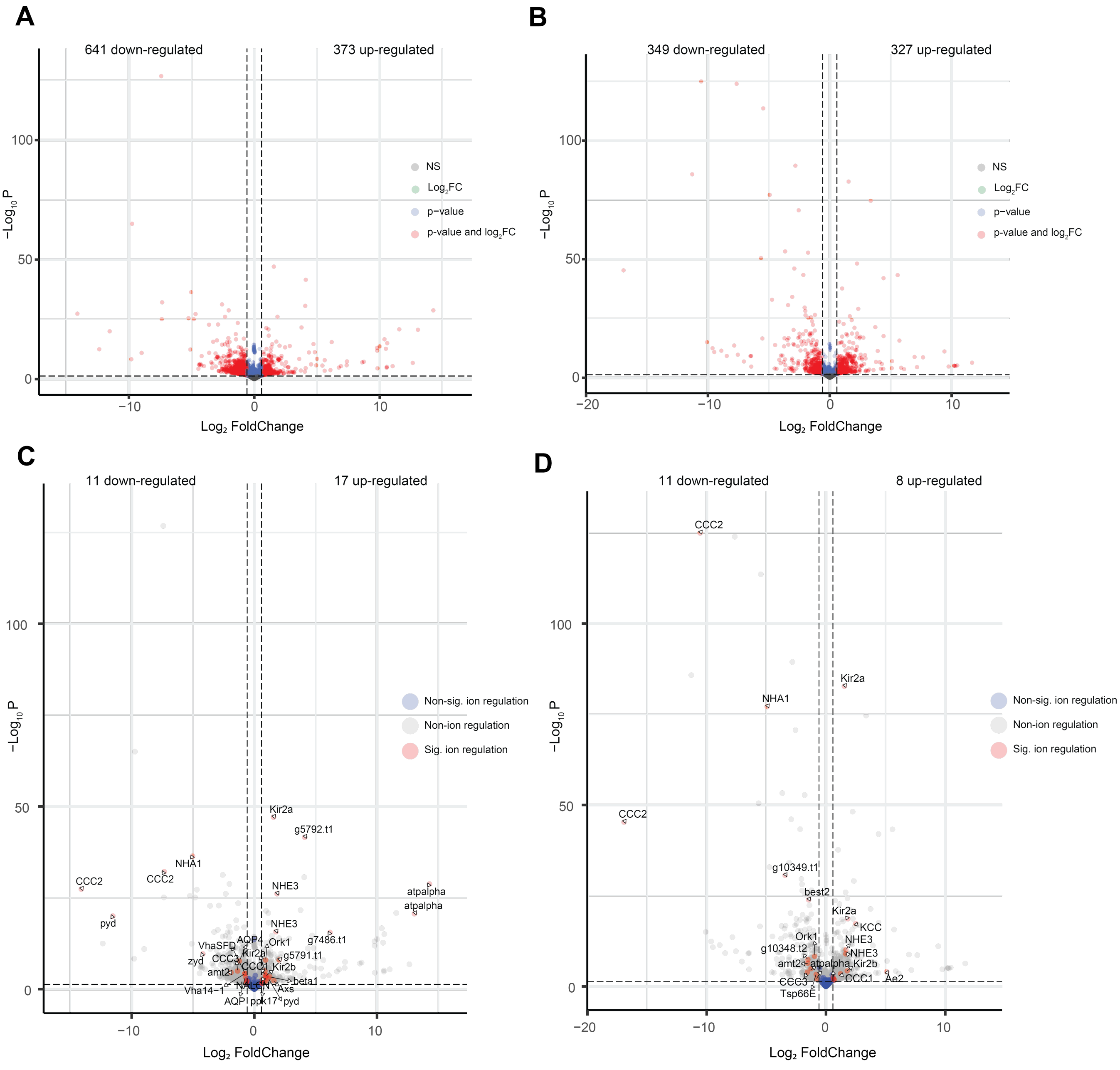


**Supplementary Figure 6. Differentially expressed transcripts between FW and SW conditions in whole larval and anal canal datasets.**
Differential expression between FW and SW conditions in (A) *Ae. togoi* whole larvae transcripts, (B) *Ae. togoi* larval anal canals transcripts, (C) ion regulatory transcripts from whole *Ae. togoi* larvae, and (D) ion regulatory transcripts from *Ae. togoi* larval anal canals. (A) 1,014 whole larval transcripts were differentially expressed out of 18,391 total transcripts. (B) 886 anal canal transcripts were differentially expresssed out of 18,391 total transcripts (C) 28 whole larval ion regulatory transcripts were differentially expressed out of 314 ion regulatory transcripts. (D) 19 anal canal ion regulatory transcripts were differentially expressed out of 314 ion regulatory transcripts. (A-D) Significance was determined by a threshold cutoff of p-value > 0.05 and $Log_2$FoldChange > $Log_2(1.5)$. Plots were generated with EnhancedVolcano.
